# Memory destabilization during reconsolidation – A consequence of homeostatic plasticity?

**DOI:** 10.1101/2021.02.05.429950

**Authors:** F.E. Amorim, R.L. Chapot, T.C. Moulin, J.L.C. Lee, O.B. Amaral

## Abstract

Remembering is not a static process: when retrieved, a memory can be destabilized and become prone to modifications. This phenomenon has been demonstrated in a number of brain regions, but the neuronal mechanisms that rule memory destabilization and its boundary conditions remain elusive. Using two distinct computational models that combine Hebbian plasticity and synaptic downscaling, we show that homeostatic plasticity can function as a destabilization mechanism, accounting for behavioral results of protein synthesis inhibition upon reactivation with different reexposure times. Furthermore, by performing systematic reviews, we identify a series of overlapping molecular mechanisms between memory destabilization and synaptic downscaling, although direct experimental links between both phenomena remain scarce. In light of these results, we propose a theoretical framework where memory destabilization can emerge as an epiphenomenon of homeostatic adaptations prompted by memory retrieval.

## INTRODUCTION

After retrieval, processes such as reconsolidation or extinction can alter the content and/or strength of memories. Behavioral studies typically describe these two phenomena as separate entities that are triggered by distinct retrieval conditions. In fear conditioning, for example, a long session of nonreinforced reexposure to the context leads to a decrease in the conditioned response through extinction, while shorter reexposure durations lead to reconsolidation and reinstate susceptibility to amnestic agents (Bustos et al., 2009; Pedreira and Maldonado, 2003). Other boundary conditions such as the strength and age of the memory and the degree of prediction error can also influence which process will occur (Eisenberg et al., 2003). However, it is still unclear how the transition between these opposite behavioral outcomes develops (Almeida-Corrêa and Amaral, 2014; Cassini et al., 2017; Merlo et al., 2014).

Pharmacological interventions such as protein synthesis inhibitors can block both reconsolidation and extinction, suggesting that similar plasticity systems might underlie the two phenomena. To address this hypothesis, Osan et al. (2011) investigated the transition between them using an attractor neural network model, in which the interaction between Hebbian plasticity and a mismatch-driven synaptic weakening process led to different behavioral outcomes, depending on the similarity between the original learning and the new experience. Although their results showed that both reconsolidation and extinction could be produced by similar plasticity systems, the mismatch-induced degradation term was not related to a biologically plausible form of plasticity, making this a largely theoretical proposal.

Nevertheless, pharmacological evidence does suggest the existence of a memory destabilization system, which includes mechanisms such as NMDA receptors (Ben Mamou et al., 2006; Milton et al., 2013; Nakayama et al., 2016; Yu et al., 2016), the ubiquitin-proteasome system (Da Silva et al., 2013; Fustiñana et al., 2014; Lee, 2008; Lee et al., 2008), CB1 receptors (Lee et al., 2019), L-type voltage-gated calcium channels (LVGCCs) (Suzuki et al., 2008) and calcineurin (Fukushima et al., 2014; Yu et al., 2016). This set of mechanisms, derived from behavioral studies in which destabilization is blocked by pharmacological agents, can be useful to speculate on possible synaptic correlates of memory destabilization.

Interestingly, there is evidence that some forms of negative synaptic plasticity share molecular mechanisms with memory labilization. The induction of long-term depression (LTD), for example, requires NMDA receptors in the hippocampus (Dudek and Bear, 1992) and involves endocannabinoids as retrograde messengers in the striatum (Gerdeman et al., 2002), neocortex (Nevian and Sakmann, 2006) and cerebellum (Qiu and Knöpfel, 2009). Beyond Hebbian mechanisms, synaptic weakening can also be induced by homeostatic plasticity, which adjusts synapse number and/or strength in response to chronic changes in neural activity (Marder and Goaillard, 2006; Turrigiano, 2012). Several types of plasticity can drive homeostatic adjustments at the cellular or network level (**Table 1**). Among these, synaptic weakening can occur through synaptic scaling (Turrigiano, 2008), heterosynaptic plasticity (Chistiakova et al., 2014), and sliding threshold modifications (Keck et al., 2017).

**Table 1:** Homeostatic plasticity types and their features. *Spatial scale* refers to which spatial extent a given plasticity operates. Local mechanisms act on individual or small groups of synapses within a neuron, while other processes are able to evoke cell-wide or circuit-wide responses for excitability adjustment. *Onset time* denotes the time range in which each type of plasticity is commonly observed. *Spike-time dependence* describes whether a form of plasticity follows spike-timing plasticity rules. *NMDA dependence* describes whether it depends on NMDA receptors. *Neuronal response to activity change* briefly describes the circuit and cellular adaptations to chronic changes in the firing rate. BCM, Bienenstock-Cooper-Munro; NMDAR, N-methyl-D-aspartate receptor; LTP, long-term potentiation; LTD, long-term depression.

| Homeostatic Plasticity type | Spatial scale | Onset time | Spike-time dependence | NMDA dependence | Response mechanisms | References |
| --- | --- | --- | --- | --- | --- | --- |
| **Synaptic scaling** | Global (cell-wide) | Hours to days | Downscaling only | Downscaling only | Multiplicative change in synaptic weights | (Siddoway et al., 2014; Turrigiano, 2008) |
| **Heterosynaptic plasticity** | Local synapses | Minutes to hours | Yes | Yes | Average changes in the weights of specific synapses, depending on initial synaptic properties | (Chistiakova et al., 2015, 2014) |
| **Sliding threshold (BMC model)** | Global (cell-wide) | Days | Yes | Yes | LTP/LTD threshold change depending on initial synaptic properties | (Bridi et al., 2018; Tara Keck et al., 2017) |
| **Excitation/inhibition balance** | Global (circuit-wide) | Hours to days | No | Yes | Circuit-wide remodeling of inhibitory synapse clusters and dendritic spines | (Chen et al., 2012; Sohal and Rubenstein, 2019) |
| **Intrinsic excitability modulation** | Local or cell-wide | Seconds to hours | No | Yes | Modifications of membrane conductance and voltage-gated ion channel expression, either locally or cell-wide. | (Debanne et al., 2019; Marder and Goaillard, 2006) |

Unlike Hebbian plasticity, which is typically fast, homeostatic plasticity can take place across a variety of timescales (Zenke and Gerstner, 2017). Synaptic downscaling in response to increased neuronal firing, for example, can take several hours to occur (Turrigiano et al., 1998), matching the relatively slow decay in conditioned responses (Nader et al., 2000) and in synaptic potentials (Fonseca et al., 2006) after protein synthesis inhibition upon memory reactivation. Moreover, the activity dependence of synaptic downscaling makes it an interesting candidate to explain why memory destabilization requires reactivation to occur. Finally, synaptic downscaling also requires mechanisms implicated in labilization, such as NMDA receptor activation (Lee and Chung, 2014), AMPA receptor endocytosis (Shepherd et al., 2006), LVGCC activity (Goold and Nicoll, 2010; Lee and Chung, 2014), and protein degradation (Jakawich et al., 2010).

To investigate whether homeostatic plasticity could be a feasible mechanism for retrieval-induced memory destabilization, we performed a computational study using two different network models, previously developed to study each of the processes separately (Osan et al., 2011; Auth et al., 2020). This was followed by systematic reviews of the molecular mechanisms of both phenomena and of existing links between homeostatic plasticity and memory destabilization in the literature.

## RESULTS

In order to study whether memory destabilization could arise from homeostatic plasticity, we used adaptations of two previously published computational models. The first (Osan et al., 2011) studied the transition between reconsolidation and extinction in an attractor network, with labilization driven by mismatch between the training and reactivation patterns. The second (Auth et al., 2020), used a combination of Hebbian plasticity and homeostatic synaptic scaling to allocate stimuli as internal representations in a memory network (Auth et al., 2020).

In both models, we focus on whether synaptic scaling can mediate the different effects of Hebbian plasticity blockade on stored memories under various reexposure conditions. More specifically, we seek to model the general results observed in Suzuki et al., (2004), in which a transition from simple retrieval to reconsolidation to extinction is observed with increasing reexposure duration, leading to different effects of protein synthesis inhibition in each case. As in Osan et al. (2011), we assume that longer reexposure sessions are associated with increasing mismatch between the training and reexposure context representations, and model the patterns accordingly in both models.

### Investigating homeostatic plasticity as a destabilization mechanism in an attractor model of reconsolidation and extinction

Our first model is an adaptation of the attractor network described in Osan et al. (2011). This fully connected Hopfield-like network is composed of 100 neurons exhibiting graded activity from 0 to 1, driven by their recurrent connections and by an input cue that simulates an animal’s current representation of its environment (**Figure 1a**). In the original model, changes in recurrent connection weights occurred through a combination of Hebbian plasticity and a mismatch-induced degradation term, which weakened synapses causing mismatch between the input cue and the pattern retrieved by the network. The combination of these two forms of plasticity led to transitions between simple retrieval, reconsolidation and extinction according to the degree of mismatch between initial learning and reexposure, used by the authors to model the duration of contextual reexposure in fear conditioning.

**Figure 1:**
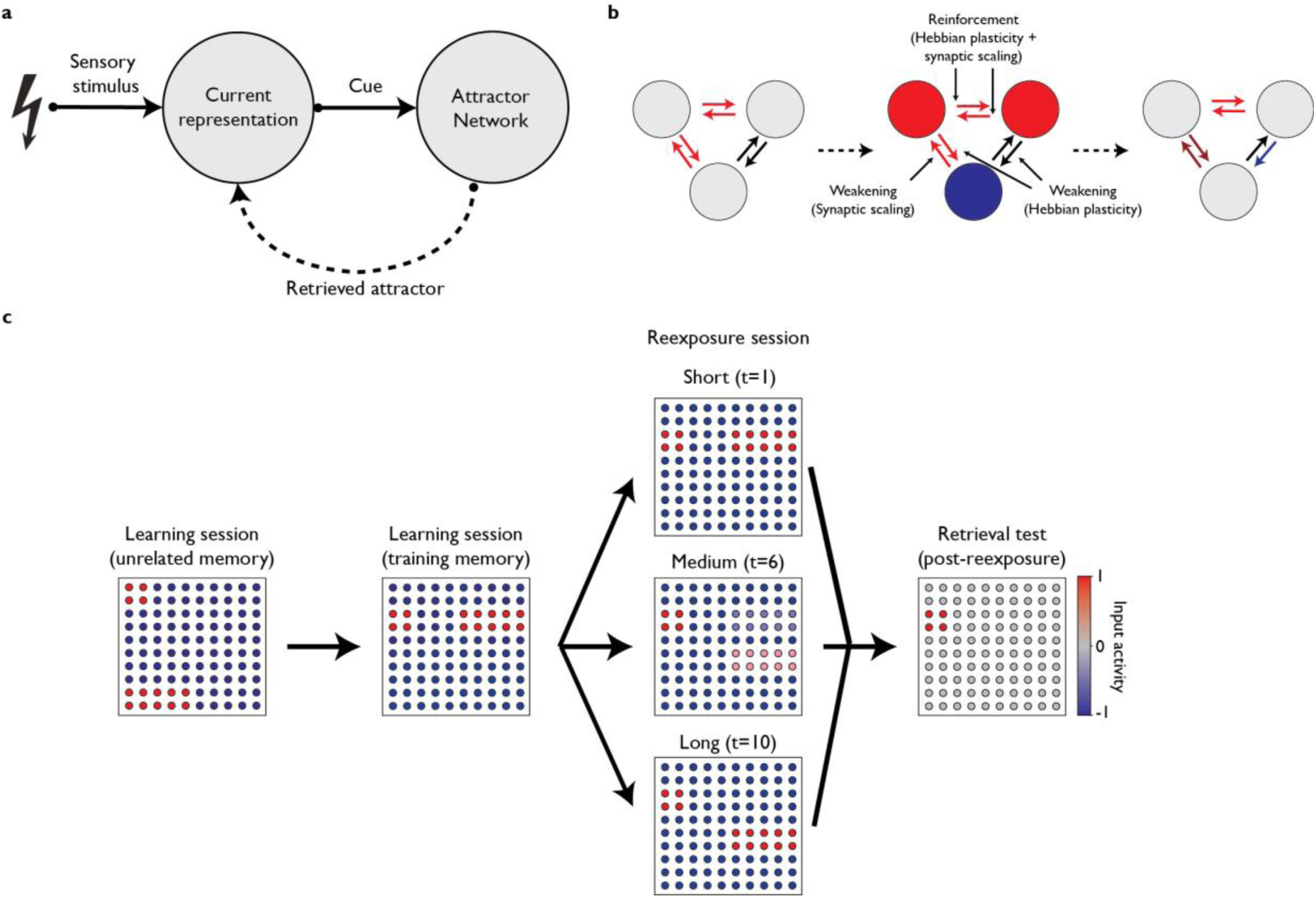
Attractor network model adapted from Osan et al. (2011) **(a)** Network architecture. The model is composed of two networks, representing an animal’s current content representation and a memory storage area. Connections from the memory network to the current representation are assumed but not explicitly modeled. **(b)** Network plasticity mechanisms. Simultaneous activation of two neurons (red) leads them to reinforce their connections through Hebbian plasticity. The Hebbian plasticity term also leads to formation of inhibitory connections from active to inactive neurons (blue), presumably mediated by inhibitory neurons. Increased activity in a neuron leads to activation of the synaptic scaling term, which lowers the neuron’s input weights globally, leading to weakening of synapses when Hebbian reinforcement is absent. **(c)** Modeling of fear conditioning and reactivation. After learning an unrelated pattern and the training memory, the network undergoes a reexposure session with an input of varying similarity to the training pattern according to reactivation duration, with longer durations leading to progressively dissimilar patterns. After this, retrieval is measured by activating the shared neurons between both patterns (i.e. the context) and observing the attractor to which the network evolves.

In our adaptation of the model, the mismatch-degradation term is replaced by a synaptic scaling term, which adjusts a neuron’s synaptic input weights according to its firing rate. This term, adapted from Tetzlaff et al. (2013), compares a neuron’s firing rate to a desired target activity, weakening all excitatory connections received by the neuron if this target is exceeded. Thus, while Hebbian plasticity reinforces the connections between two active neurons, synaptic weakening due to scaling occurs when the postsynaptic neuron is highly active in the absence of presynaptic activity (**Figure 1b**).

The Hebbian learning term is associated with a parameter S that represents the biochemical requirements for this form of plasticity. As described in Osan et al. (2011), we vary S to simulate the effects of protein synthesis inhibitors or plasticity-enhancing drugs in consolidation, reconsolidation, and extinction. Similarly, the synaptic scaling term is associated with a parameter κ that represents the ratio between Hebbian plasticity and synaptic scaling. Variation in κ is used to simulate the effects of destabilization blockers or enhancers in reconsolidation.

We model conditioning and reexposure to a context using a sequence of three different cue patterns (**Figure 1c**). After learning a cue pattern unrelated to fear conditioning, the network receives a training cue activating two groups of neurons, one representing the context (4 neurons) and another representing the fearful experience (10 neurons), leading to reinforcement of their connections through Hebbian plasticity. The third cue pattern simulates a reexposure session, with activation of context neurons alongside a variable mix of fear neurons and a separate group of 10 neurons representing a safe environment. Short reexposure sessions are modeled by learning cues that are similar to the training pattern, assuming that presentation of the context will initially trigger a strong fear response. For longer reexposure sessions, activation of these neurons is gradually replaced by that of safety neurons as a function of reexposure time (t), until the extinction pattern (with full activation of the safety neurons and no activation of fear neurons) is reached at the maximum duration (t=10). The retrieval test is simulated by presenting a cue pattern activating the context neurons and observing the activation pattern to which the memory network evolved. As in the original model, retrieval of the training memory attractor is assumed to lead to greater freezing behavior than other attractors (see Methods for more detail).

Results obtained with the model are shown in **Figure 2**. Under normal conditions, the memory network is able to form and retrieve the association between context and fear neurons, leading to retrieval of this memory and high freezing at the cued retrieval test. Blockade of Hebbian plasticity during the training session prevents this process, leading to a decrease in test freezing (**Figure 2a**), as observed with protein synthesis inhibitor (PSI) administration before or shortly after fear conditioning (Schafe et al., 1999; Schafe and LeDoux, 2000).

**Figure 2:**
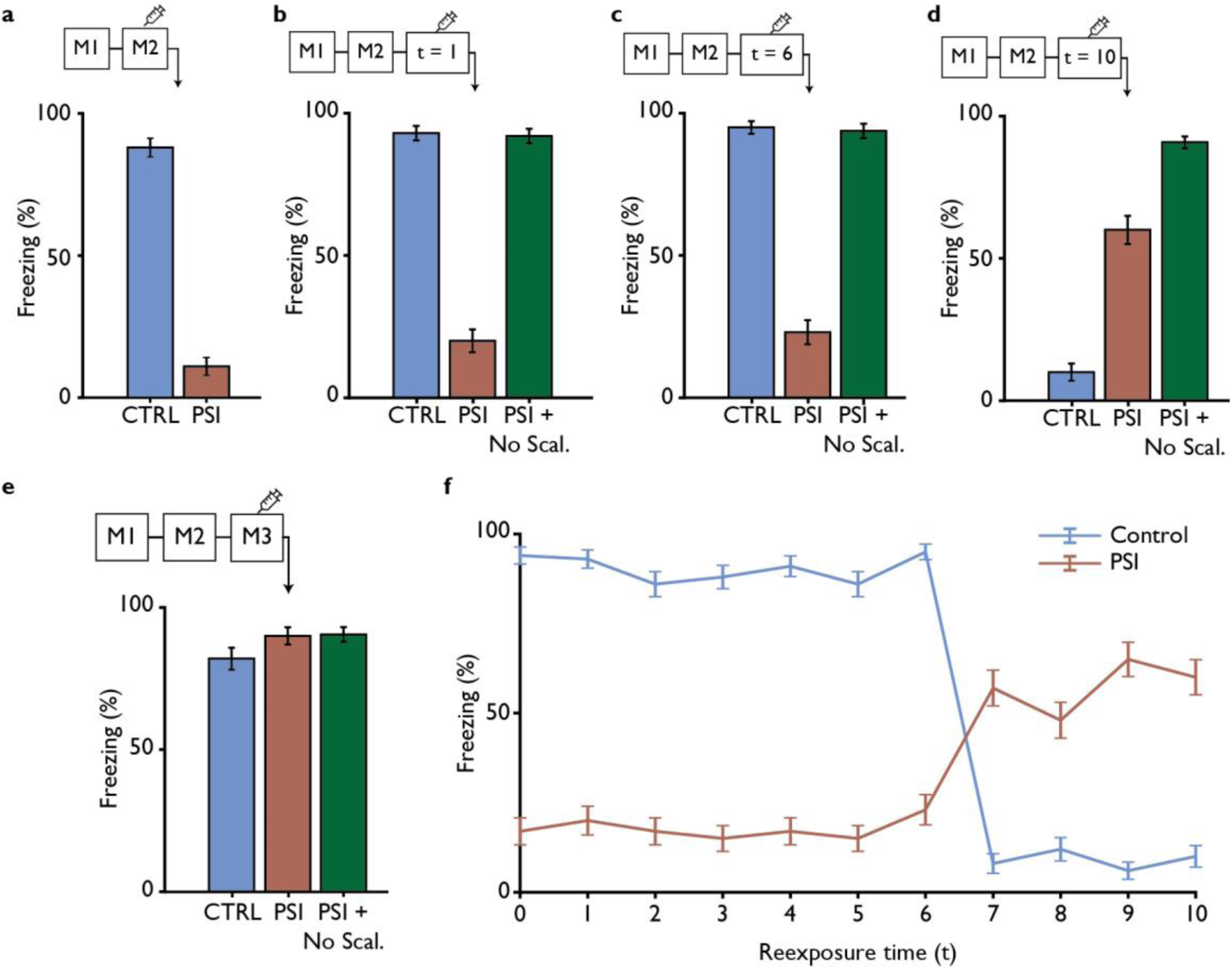
Effects of Hebbian plasticity blockade in different protocols. Bars represent mean ± SEM of freezing values in the retrieval test, calculated on the basis of the retrieved attractor (see Methods), for 100 simulations with control conditions (red), Hebbian plasticity blockade by PSI injection (blue) or simultaneous Hebbian plasticity and synaptic scaling blockade (green). Treatments occur at different moments, as shown by the timelines above graphs, in which M1 representing an unrelated pattern, M2 corrsponds to the training memory and t indicates the reexposure duration. **(a)** Amnestic effect of Hebbian plasticity blockade (PSI) during initial learning of the shock memory. **(b)** Amnestic effect of Hebbian plasticity blockade in a short (t=1) reexposure session. **(c)** Amnestic effect of Hebbian plasticity blockade in an intermediate (t=6) reexposure session. **(d)** Extinction disruption by Hebbian plasticity blockade in a long (t=10) reexposure. **(e)** No effect of plasticity blockade when an unrelated pattern (M3) is used as an input. Blockade of synaptic scaling (No Scal.) reverses the effect of PSIs in reconsolidation conditions (**b** and **c**) but potentiates extinction blockade in (**d). (f)** Summary of test freezing for the control and PSI groups using reexposure of varying times. A nonlinear transition between reconsolidation and extinction is observed between t=6 and t=7, leading to opposite effects of PSIs in reconsolidation and extinction.

The effect of blocking Hebbian plasticity during reactivation, meanwhile, is contingent on reexposure duration (i.e., the degree of mismatch between training and reactivation cues). Short (t=1, **Figure 2b**) or intermediate (t=6, **Figure 2c**) reexposure times lead to retrieval and reinforcement of the shock memory in the control group. In this case, blockade of Hebbian plasticity during reactivation leads to decay of the original memory, as observed experimentally with PSI injection (Nader et al., 2000). For longer reexposure durations, the control group network converges to the extinction pattern, leading to low freezing at retrieval. In this case, extinction is blocked by inhibition of Hebbian plasticity, causing freezing to be higher than in control conditions (**Figure 2d**). Blockade of synaptic scaling reverses the memory decay caused by PSIs in reconsolidation conditions, but potentiates their effect in blocking extinction.

In accordance with experimental studies (Nader et al., 2000), the effect of PSIs during reexposure does not occur if the original learning context is replaced by an unrelated cue pattern (**Figure 2e**). The effect of blocking Hebbian plasticity during low and intermediate reexposure is also abolished by blocking synaptic scaling (**Figure 2b-c**), (Ben Mamou et al., 2006; Lee et al., 2008). Parameter analysis of the model’s behavior when different values of S (**Figure S1**) or κ (**Figure S2**) are used during reexposure shows that the effects of PSIs are ‘dose-dependent’, with greater inhibition leading to larger effects, while those of synaptic scaling blockade have sharper thresholds. We also evaluate how the strength of the original learning influences reconsolidation and extinction (Eisenberg et al., 2003; Suzuki et al., 2004) (**Figure S3**). Different S values in training cause transitions between both processes to occur at different reexposure times in the control group, although very low training strength leads to low freezing at the retrieval test irrespective of reexposure time or protein synthesis inhibition.

A summary of results for varying reexposure times can be found in **Figure 2f**, in which a nonlinear transition between reconsolidation and extinction appears around t=7 with standard parameters. These results differ from those of Osan et al., 2011 in that memories are sensitive to reconsolidation even with short reexposure times (i.e., in the absence of mismatch). This occurs because the Hopfield-like framework leads to very accurate pattern completion at reexposure when the attractor is reached, leading to similar network activity for short and moderate reexposure times (**Figure S4**). Thus, plasticity rules that are based solely on the activity of the memory network (as in the case of our model) will lead to similar results between these conditions and thus fail to match experimental results in which reconsolidation blockade is contingent upon mismatch (Morris et al., 2006; Pedreira et al., 2004). Nevertheless, it is possible that mismatch dependence can occur in models with noisier retrieval, in which patterns retrieved during reexposure become progressively dissimilar from training as mismatch increases.

### Generating reconsolidation and extinction-like behavior in a network model of homeostatic plasticity

To study whether the network behavior observed in our adaptation of Osan et al.’s model could be translated to existing computational models investigating homeostatic plasticity, we used a memory allocation model adapted from Auth et al. (2020). This model was originally used to show that a combination of Hebbian learning and synaptic scaling could account for pattern separation in a recurrently connected network, allocating distinct memories to different neuronal populations when partially overlapping cue patterns were presented.

The model is composed of an input network and a memory network, as well as an inhibitory unit connected to the latter (**figure 3a**). The input area is composed of 36 rate-coded neurons that transmit information to a 900-neuron memory network through random feed-forward connections, with each neuron in the latter receiving connections from 4 neurons in the former (**Figure 3b**). The memory network is stimulated by setting the firing rate of neurons in the input area to 130 Hz for active neurons and 0 for inactive ones. It stores internal representations of the environment in its recurrent connections, with each neuron connecting to neighboring neurons within a radius of 4 units in a toroidal topology (**Figure 3c**). The inhibitory unit has bidirectional connections with all neurons in the memory area and regulates global activity.

**Figure 3:**
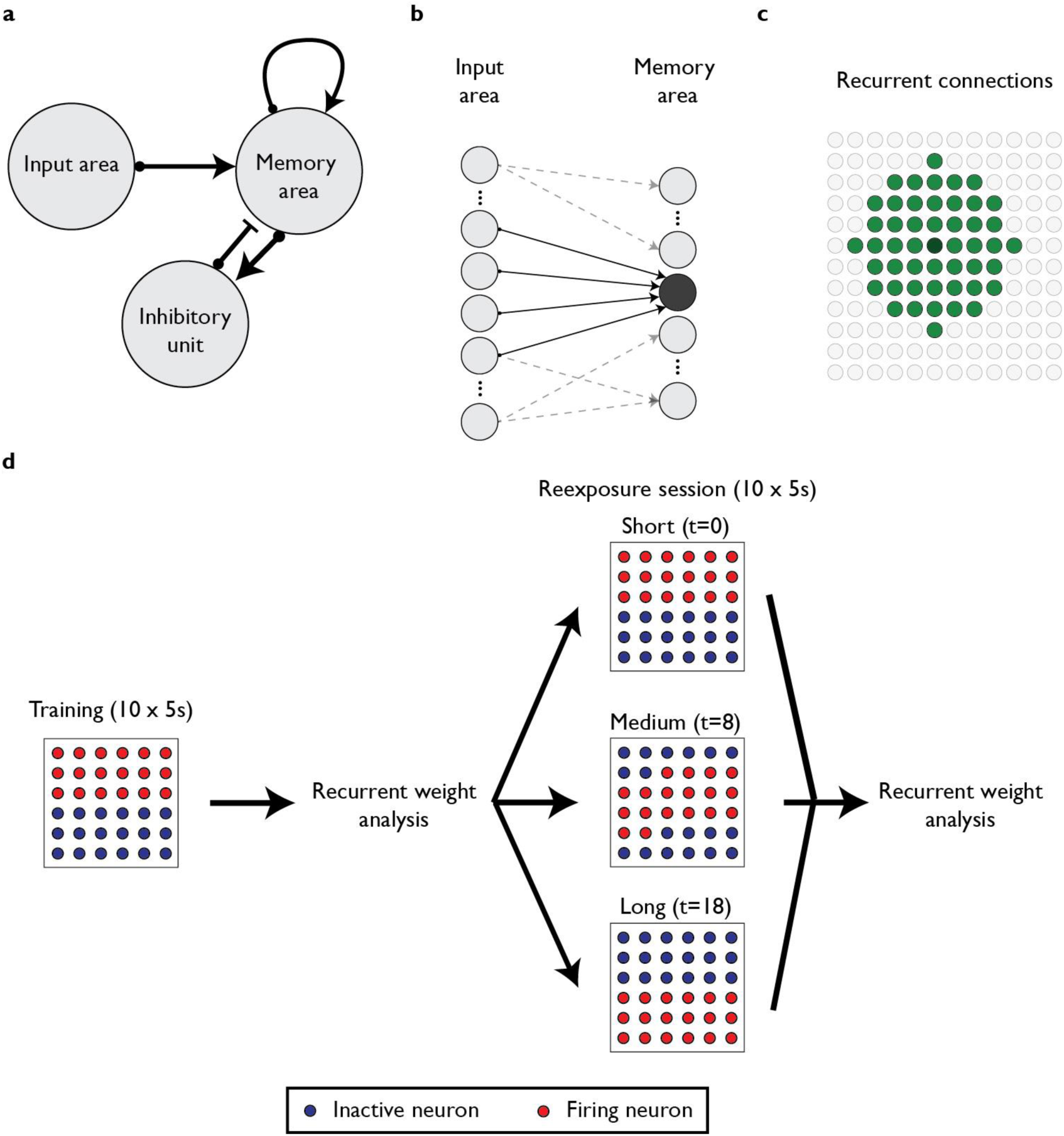
Homeostatic plasticity model adapted from Auth et al., 2020. **(a)** Context representation is encoded by the activation of the 36 neurons in the input area. These neurons stimulate the 900 neurons in the memory area, which have recurrent connections and bidirectional connections with an inhibitory unit. **(b)** Each excitatory neuron in the memory area receives connections from 4 different excitatory neurons in the input area. **(c)** Recurrent connections within the memory area are made with neighboring neurons in a radius of 4, in a toroidal topology. **(d)** Protocol used to model contextual fear conditioning with different reexposure durations. For the training session, a pattern representing the aversive stimulus is presented. Afterwards, reactivation is modeled by an input that varies according to reexposure duration. Short reexposure durations correspond to patterns similar to that of the training session, while progressively longer durations lead to gradual replacement of neurons in the original input by new ones. After each session, the recurrent weight of neurons representing each pattern is obtained by finding clusters of at least 30 neurons with mean recurrent weights above 40 (see Methods). When no cluster is found, the mean weight is considered to be equal to the global mean recurrent weight of the memory area.

We consider all connections between excitatory neurons in the model to be plastic. A Hebbian plasticity term is activated when pre- and post-synaptic neurons fire simultaneously; in our adaptation, we vary its constant *S* to simulate the effect of protein synthesis inhibitors. As in our first model, the synaptic scaling term in the model is dependent on a comparison between postsynaptic firing rate and an activity target (Tetzlaff et al.,2011; Tetzlaff et al., 2013). To simulate the effects of blocking synaptic scaling, we vary the constant *κ*, representing the ratio of Hebbian plasticity and synaptic scaling.

As in the first model, our stimulation protocol aims to model fear conditioning followed by different durations of reexposure to the context. Training consists of presenting a cue stimulus similar to those used in the original model. This is followed by a reexposure session, in which we again assume that increasing duration leads to a gradual increase in mismatch between the training and reexposure patterns. With a short reexposure time (t = 0), the same neurons from the training session are activated in the input network, while progressively longer reexposure lead to stepwise substitution of these neurons by new neurons until the pattern is completely different (i.e., a pure extinction pattern) (**Figure 3d**). Even though input cues are completely different, some degree of activation overlap by the cue can still occur in the memory area due to the random divergent connections between neurons in both networks.

After the training and reexposure sessions, we analyze the synaptic weight matrices in the memory network to look for neuronal clusters representing each memory. The fear memory cluster formed during the training session is observed both after training and after reexposure of any duration in the control group (**Figure 4a**). If there is some degree of mismatch between the training and reactivation sessions, this cluster is sometimes updated and allocated in a new region that partially overlaps with the original one. When longer reexposure times are used, a second memory cluster is formed, which is analogous to an extinction memory and coexists with the original one in the network.

**Figure 4:**
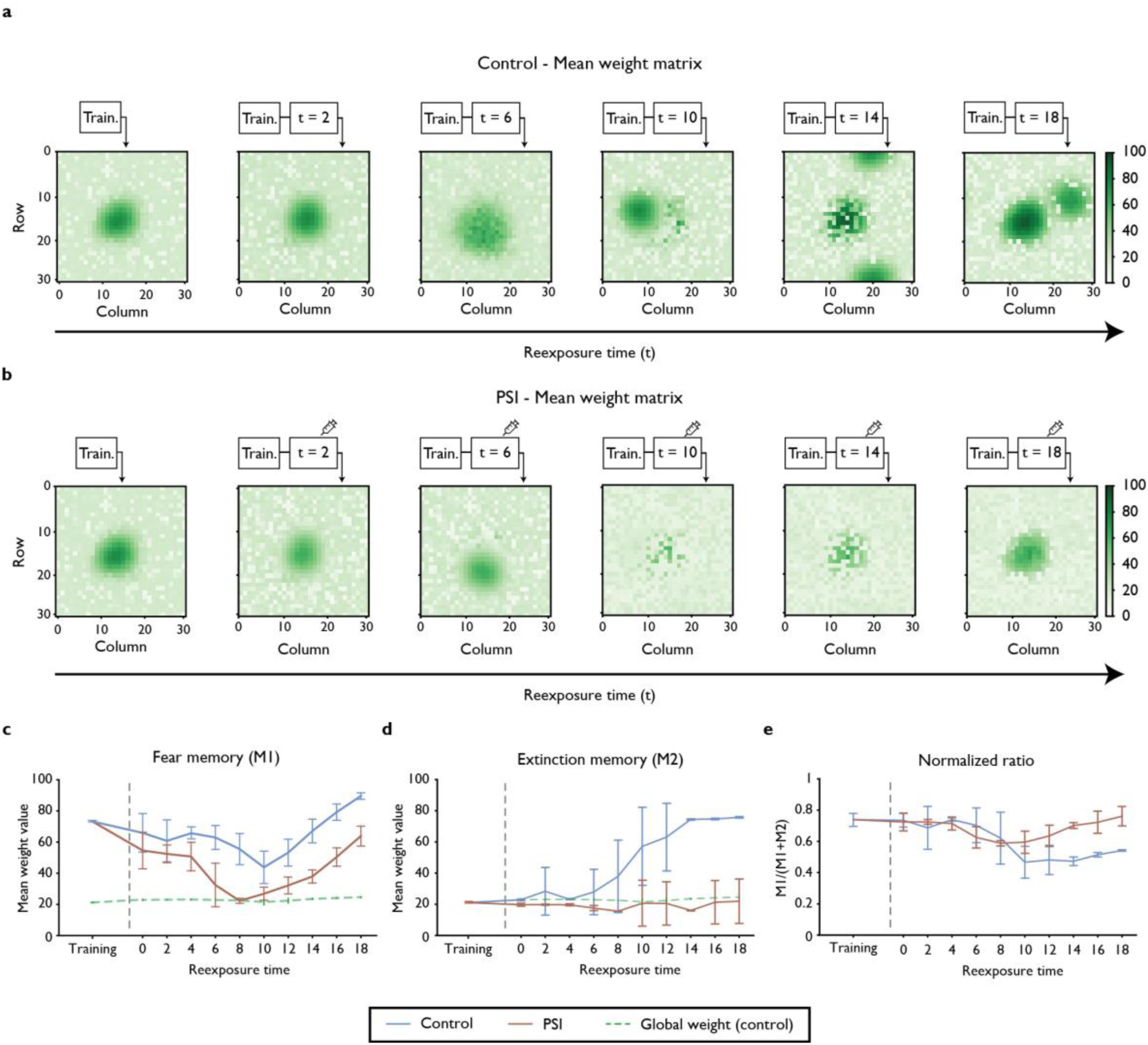
Effects of Hebbian plasticity blockade on synaptic weights within memory clusters. **(a)** Representation of recurrent synaptic weights in the memory area for the control group after training (left panel) and after reactivation with different reexposure durations in a representative simulation. Heat map represents the mean of recurrent weights for each neuron in the memory area, displayed on a 30 x 30 grid. A single cluster, corresponding to the fear memory, is observed after training and after short reexposures, while longer reexposure durations lead to the formation of a second cluster representing extinction. **(b)** Same as (a), but with protein synthesis inhibition during reexposure. Decrease of Hebbian plasticity weakens synaptic weights in the fear memory cluster, particularly in intermediate reexposure durations, and prevents the formation of the extinction cluster. **(c)** Mean weight value of the fear memory cluster in control (blue) and PSI (red) condition after training and reexposure with varying duration. Lines represent means of 20 simulations with different connection topologies and starting conditions. Dashed green line shows the global connection weight of the network under control conditions, including neurons inside and outside the clusters **(d)** Mean weight value of the extinction memory cluster after different durations of reexposure under control and PSI conditions. **(e)** Ratio between the fear and extinction memories for reexposures of different durations with and without PSIs. Hebbian plasticity blockade has little effect after short reexposure duration, decreases the ratio for intermediate durations and increases it for long durations.

When Hebbian plasticity is decreased to simulate the effect of protein synthesis inhibition, the fear memory cluster is maintained when short reexposure times are used (**Figure 4b**). With intermediate reexposure durations, however, the decrease of Hebbian plasticity leads to marked weakening of synaptic weights in the original memory cluster, mediated by uncompensated synaptic scaling. With longer reexposure times, the fear memory is preserved, but formation of the extinction cluster is blocked. These results are qualitatively similar to the ones obtained in the previous model, with the exception that mismatch is now required for reconsolidation blockade, as occurs in Osan et al. (2011) and in several experimental studies (e.g. Morris et al., 2006; Pedreira et al., 2004).

**Figure 4c** shows the mean recurrent weight of the cluster representing the training session for simulations with different reexposure durations. Protein synthesis inhibitors lead to a reduction in synaptic strength for all durations, but this effect is more marked in intermediate reexposure times (i.e., t values between 6 and 8), and is reduced for longer reexposure sessions in which extinction is observed. Meanwhile, the extinction memory cluster starts to appear between *t* values of 8 to 10, reaching a plateau for *t* ≥ 14, but is not formed when Hebbian plasticity is blocked (**Figure 4d**). When a normalized ratio is used to assess the balance between the fear and extinction memories (**Figure 4e**), the net effect of PSIs on this measure (which is analogous to test freezing) is neutral at short reexposure times, negative at intermediate times and positive at longer times, replicating what is observed in Osan et al. (2011) and in experimental studies using varying durations of fear conditioning (Bustos et al., 2009; Suzuki et al., 2004).

We checked the robustness of our analysis by using different minimum cluster sizes and weight thresholds for clusters. Variation in minimum cluster size (ρ) has negligible effects on the mean weight values observed (**Figure S5**), while mean weight threshold does affect the results when set too low or too high, but not for values between 30 and 50 neurons (**Figure S6**).

### Investigating molecular mechanisms of memory destabilization in the literature

To investigate whether experimental data support the possibility that homeostatic plasticity underlies memory destabilization, we performed a systematic review to investigate the molecular mechanisms implied in behavioral studies of destabilization (see Methods for search terms and other details). After screening 769 articles, we extracted a total of 88 experiments from 41 studies that investigated the effect of a pharmacological or genetic manipulation on reconsolidation blockade caused by another intervention (**Figure S7**). The molecular targets analyzed in these studies are presented according to brain structure on **Table 2**.

**Table 2:**
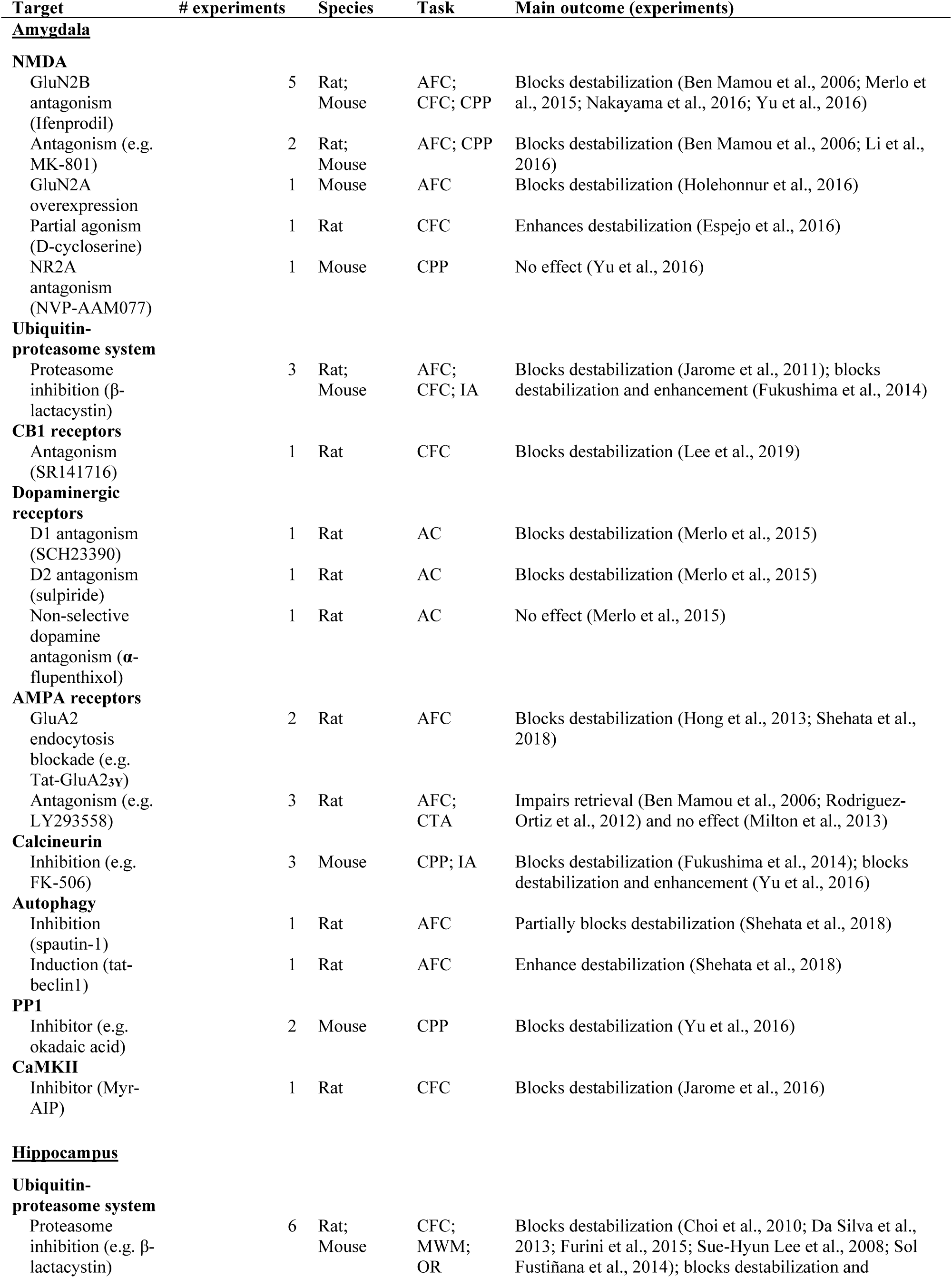
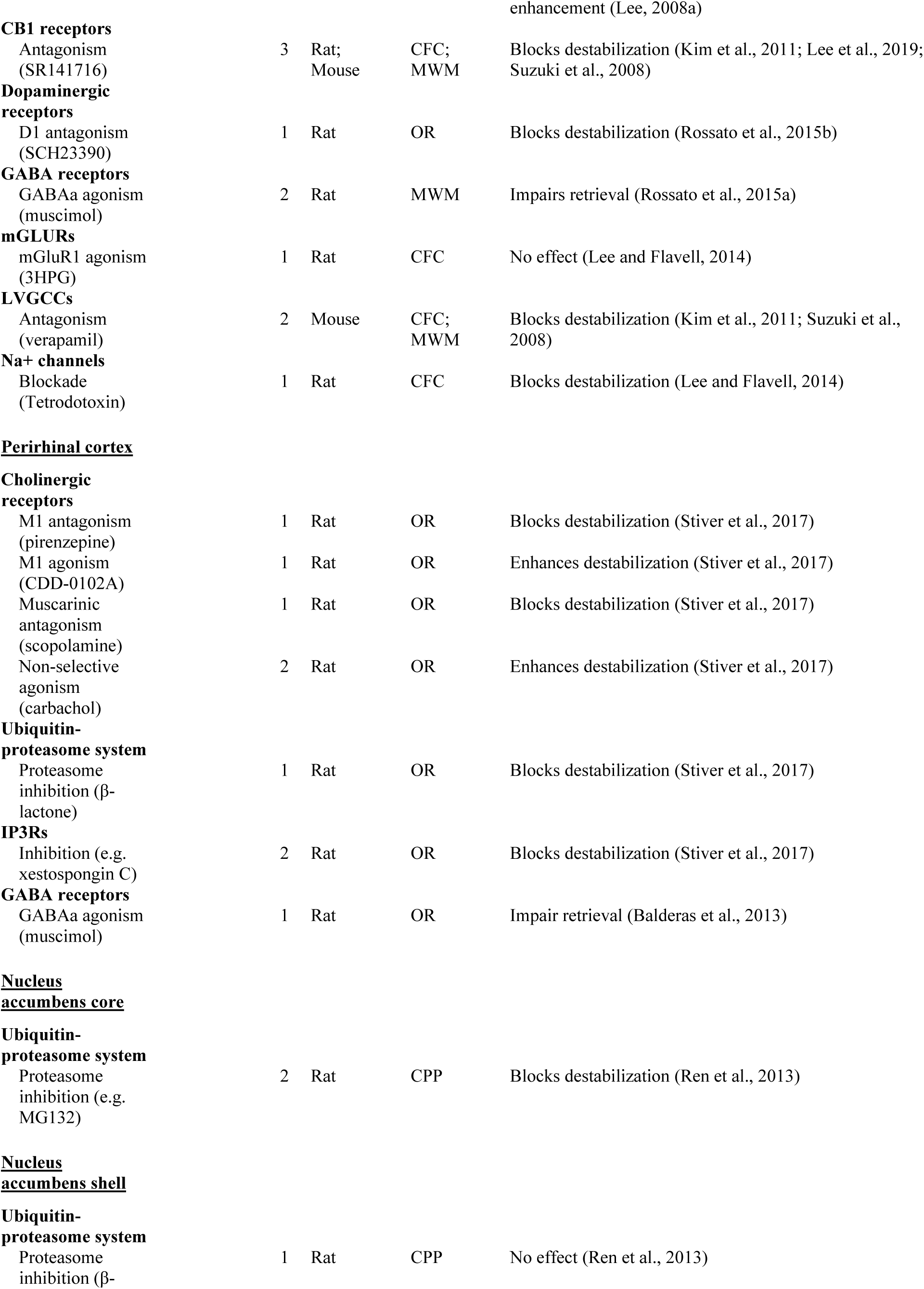
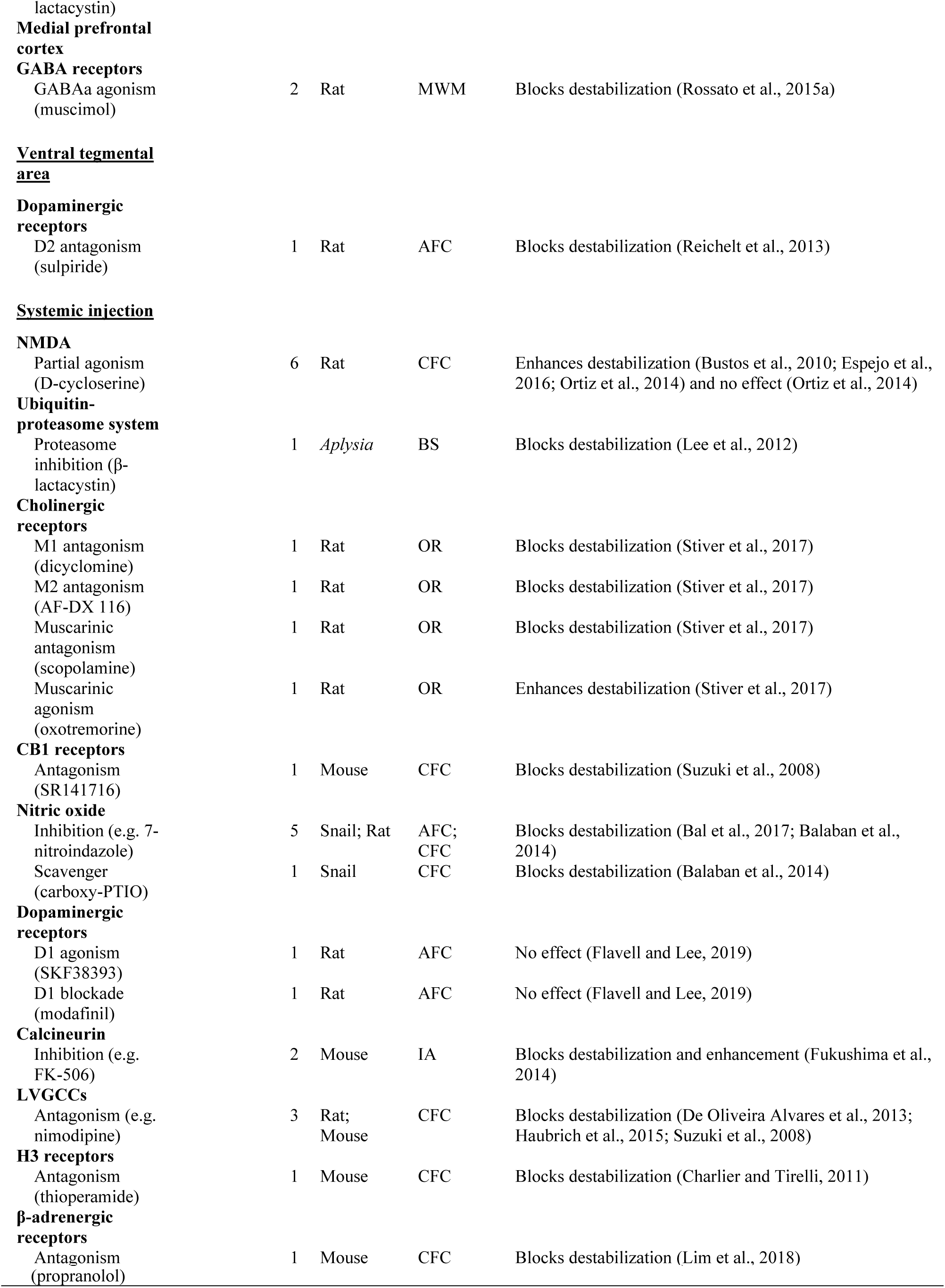
Molecular mechanisms for memory destabilization. AC, appetitive conditioning; AFC, auditory fear conditioning; AMPA, α-amino-3-hydroxy-5-methyl-4-isoxazolepropionic acid; BS, behavioral sensitization; CaMKII, Calcium/calmodulin-dependent protein kinase type II; CB1, endocannabinoid receptor type 1; CFC, contextual fear conditioning; conditioned place preference; CPP, conditioned place preference; CTA, conditioned taste aversion; GABA, γ-aminobutyric acid; H3, histamine H3 receptors; IA, inhibitory avoidance; IP3Rs, inositol trisphosphate receptors; LVGCCs, L-type voltage-gated calcium channels; MWM, Morris water maze; NMDA, N-methyl D-aspartate receptor; OR, object recognition; PP1, protein phosphatase 1;

Most structure-specific studies were aimed at the amygdala or hippocampus, while others targeted the whole brain through systemic injections. For intra-amygdala infusions, NMDA receptors were the most commonly studied mechanism, with both nonspecific and GluN2B antagonists shown to block destabilization. In the hippocampus, the ubiquitin-proteasome system was the most studied mechanism, with its inhibition found to block not only destabilization, but also memory enhancement in some studies. LVGCCs and CB1 receptors were also shown to block destabilization in the hippocampus in multiple studies. CB1 and dopaminergic receptors, as well as the ubiquitin-proteasome system, were implicated in memory destabilization in the two structures, suggest that a similar labilization system could be at work in both brain regions.

Other brain regions studied were the perirhinal cortex, nucleus accumbens, medial prefrontal cortex and ventral tegmental area, but these were evaluated in isolated studies investigating particular mechanisms. Studies with systemic injections in rodents confirmed the role of NMDA receptors, LVGCCs and CB1 receptors, and added other candidates such as nitric oxide and muscarinic, β-adrenergic and H3 receptors, although these were evaluated in a smaller number of studies. Interestingly, studies in *Aplysia* also found the ubiquitin-proteasome system and nitric oxide to be important for destabilization, suggesting that this memory labilization system is present in invertebrates as well.

Of note, experiments with negative results – i.e., showing the lack of effect of a pharmacological manipulation on destabilization – were very uncommon, comprising only 9% of our sample (this does not include studies of boundary conditions that show a negative effect of an intervention in a particular condition – i.e., dose or timing – but a positive one in another, which are shown on **Table S1**). Moreover, most of these were drawn from studies in which another intervention was found to be effective. This suggests that publication bias is likely in this field of study, meaning that the apparent consistency in the results included in the table should be interpreted with caution. Lack of publication of negative results also makes it harder to evaluate whether discrepancies in labilization mechanisms exist across structures or species.

### Involvement of destabilization mechanisms in synaptic downscaling

To investigate the relationship between the molecular mechanisms underlying memory destabilization and homeostatic plasticity, we went on to review which of the components described in Table 2 have been studied in experimental models of homeostatic plasticity. For this, we focused on synaptic downscaling, which is arguably the best studied model of homeostatic plasticity, and corresponds more closely to the implementation used in our models. By combining data from a synaptic scaling systematic review dataset (Moulin et al., 2020) and manual searches in PubMed (see Methods), we identified 13 articles that conducted experiments studying 9 molecular mechanisms of memory destabilization (**Table 3**). Importantly, this search strategy focused on identifying whether established mechanisms of destabilization are also important for downscaling; thus, it would not be expected to identify mechanisms of downscaling that are not involved in destabilization.

**Table 3.** Role of labilization mechanisms in synaptic downscaling. Columns summarize the main findings, models and intervention methods of articles investigating the role of memory destabilization components in synaptic downscaling. AMPAR, α-amino-3-hydroxy-5-methyl-4-isoxazolepropionic acid; APV, D(−)-2-Amino-5-phosphonopentanoic acid; CaMKII, Calcium/calmodulin-dependent protein kinase type II; ChR2, channelrhodopsin-2; UV, ultraviolet; DRB, 5,6-dichloro-1-beta-D-ribobenzimidazole. hipp, hippocampal; K.O., knockout; LVGCC, L-type voltage-gated calcium channels; MeCP2, methyl-CpG binding protein 2; NMDA, N-methyl D-aspartate receptor; PP1, protein phosphatase 1; TTX, tetrodotoxin.

| Mechanism | Interference method | Excitation mechanism | Biological model | Effect on downscaling |
| --- | --- | --- | --- | --- |
| **NMDA Receptors** | 1. APV<br>2. APV/ ifenprodil<br>3. APV<br>4. APV | 1. Optogenetics (ChR2)<br>2. Light-gated GluR6 stimulation<br>3. Bicuculline<br>4. Bicuculline | 1. Mice organotypic hipp. cultures.<br>2. Rat primary hipp. cultures.<br>3. Rodent primary cortical cultures and hipp. slices<br>4. Rat primary hipp. cultures | 1. No effect (Goold and Nicoll, 2010)<br>2. Inhibition of AMPAR single-synapse downscaling (Hou et al., 2011)<br>3. Inhibition of AMPAR downscaling (Qiu et al., 2012)<br>4. Inhibition of downscaling (Lee and Chung, 2014) |
| **Ubiquitin-proteasome system** |  |  |  |  |
| **Proteasome function** | Lactacystin | Bicuculline | Rat primary hipp. cultures | Inhibition of downscaling (Jakawich et al., 2010) |
| **Nedd4 (E3 ubiquitin ligase)** | 1. N.A.<br>2. Nedd4-1 shRNA hairpin expression<br>3. Lentiviral-induced expression of Nedd4-2 | 1. Light-gated GluR6 stimulation<br>2. Bicuculline<br>3. Picrotoxin | 1. Rat primary hipp. cultures.<br>2. Rat primary hipp. and cortical cultures<br>3. Primary hipp. and cortical cultures of Fmr1 KO mice | 1. Chronic synaptic stimulation increases Nedd4-1 expression (Hou et al., 2011)<br>2. Inhibition of downscaling (Scudder et al., 2014)<br>3. Rescue of downscaling in Fmr1 KO neurons (Lee et al., 2018) |
| **AMPA receptors** |  |  |  |  |
| **GluA2 expression** | Genetic deletion | Optogenetics (ChR2) | Mice organotypic hipp. cultures. | Inhibition of AMPAR downscaling (Goold and Nicoll, 2010) |
| **MeCP2 (Regulates GluA2 expression)** | 1. Genetic deletion or shRNA expression<br>2. Genetic deletion | 1. Bicuculline<br>2. Bicuculline | 1. Rat/mouse cortical primary cultures and hipp. slices<br>2. Mice hipp. primary cultures | 1. Inhibition of downscaling (Qiu et al., 2012)<br>2. Inhibition of AMPAR downscaling (Xu and Pozzo-Miller, 2017) |
| **Arc/Arg3.1 (Induces AMPAR endocytosis)** | Genetic deletion | Bicuculline | Rat/mouse hippocampal and cortical neurons. | Inhibition of AMPAR downscaling (Shepherd et al., 2006) |
| **Calcineurin** | 1. FK-506<br>2. N.A. | 1. Light-gated glut. receptor (GluR6)<br>2. Bicuculline | 1. Rat primary hipp. cultures.<br>2. Mice cortical primary cultures | 1. No effect on AMPAR downscaling (Hou et al., 2011)<br>2. Chronic stimulation reduces calcineurin levels (Diering et al., 2014) |
| **LVGCCs** | 1. Nifedipine<br>2. Nimodipine<br>3. Nifedipine<br>4. Nifedipine | 1. Optogenetics (ChR2)<br>2. Bicuculline<br>3. Bicuculline<br>4. Optogenetics (ChR2) | 1. Mice organotypic hipp. cultures.<br>2. Rat primary cortical cultures<br>3. Rat primary hipp. cultures<br>4. Rat hipp. slice culture. | 1. Inhibition of AMPAR and NMDAR downscaling (Goold and Nicoll, 2010)<br>2. Inhibition of AMPAR downscaling (Siddoway et al., 2013)<br>3. Inhibition of downscaling (Lee and Chung, 2014)<br>4. Inhibition of dendritic spine downscaling (Mendez et al., 2018) |
| **PP1** | Viral vector transfection | Bicuculline | Rat primary cortical cultures | Inhibition of AMPAR downscaling (Siddoway et al., 2013) |
| **CaMKII** | 1. myr-CaMKIIN or myr-AIP<br>2. KN62 | 1. Optogenetics (ChR2)<br>2. Light-gated GluR6 stimulation | 1. Mice organotypic hipp. cultures.<br>2. Rat primary hipp. cultures | 1. CaMKII expression inhibits AMPAR but not NMDAR downscaling. Inhibition of CaMKII has no effect on AMPAR or NMDAR downscaling (Goold and Nicoll, 2010)<br>2. No effect on downscaling (Hou et al., 2011) |
| **Na<sup>+</sup> channels** | 1. Tetrodotoxin<br>2. Tetrodotoxin<br>3. Tetrodotoxin<br>4. Tetrodotoxin | 1. Bicuculline<br>2. Optogenetics (ChR2)<br>3. Light-gated GluR6 stimulation<br>4. Optogenetics (ChR2) | 1. Rat auditory cortex slices<br>2. Mice organotypic hipp. cultures.<br>3. Rat primary hipp. cultures.<br>4. Rat hipp. slice culture. | 1. Inhibition of bicuculline-induced downscaling (Zhang et al., 2009)<br>2. No effect on optogenetically-induced AMPAR or NMDAR downscaling (Goold and Nicoll, 2010)<br>3. Inhibition of AMPAR downscaling (Hou et al., 2011)<br>4. No effect on dendritic spine downscaling (Mendez et al., 2018) |
| **Protein synthesis** | 1. Cycloheximide/ DRB/ anisomycin<br>2. Anisomycin | 1. Optogenetics (ChR2)<br>2. Light-gated GluR6 stimulation | 1. Mice organotypic hippocampal slice cultures.<br>2. Rat primary hipp. cultures. | 1. Inhibition of AMPAR, but not NMDAR, optogenetically-induced downscaling (Goold and Nicoll, 2010)<br>2. No effect on AMPAR downscaling (Hou et al., 2011) |

In these studies, we found consistent evidence that AMPA receptor endocytosis, LVGCCs and the ubiquitin-proteasome system, are necessary for synaptic downscaling. Experiments targeting NMDA receptors and Na^+^ channels demonstrated that they are required for downscaling induced by synaptic receptor activation (i.e., by bicuculline or UV-controlled presynaptic terminal excitation); however, their inhibition did not impact optogenetically-induced downscaling, in which excitation is independent of synapses. Concerning intracellular signaling, only one article suggested that the phosphatase PP1 is required for homeostatic plasticity. Calcineurin and CaMKII were found to be regulated during the homeostatic response to chronic excitation, but were not necessary for it to occur. Lastly, there is conflicting evidence on the role of protein synthesis, with different models yielding distinct results. While an effect of PSIs was observed on the AMPAR component of optogenetically induced downscaling, this was not the case for the NMDAR component in the same model, or for light-gated GluR6 stimulation-mediated downscaling.

## DISCUSSION

### Homeostatic plasticity can account for memory destabilization in computational models of reconsolidation and extinction

By using two different computational models, we demonstrate that synaptic scaling acting as a destabilization mechanism can account for the different effects of protein synthesis inhibition on reconsolidation and extinction. Our results are in agreement with behavioral experiments where a short nonreinforced contextual reexposure causes reconsolidation, whereas a longer reexposure duration leads to extinction, with opposite effects of PSIs in each case (Suzuki et al., 2004). In the case of the second model, it also mimics results in which some degree of mismatch between training and reexposure is necessary for reconsolidation to occur (Bustos et al., 2009; Suzuki et al., 2004).

Both models are based on abstract networks whose limitations should be noted, such as the absence of realistic topology, the use of non-spiking neurons, and an abstract concept of time. The results in both cases also critically depend on the assumption that mismatch between representations increases with greater durations of contextual reexposure, as postulated by Osan et al. (2011). Nevertheless, bearing in mind the constraints inherent to the models, our results suggest that destabilization after contextual reexposure could feasibly emerge as a byproduct of synaptic scaling-like homeostatic plasticity.

The hypothesis that homeostatic plasticity plays a role in memory phenomena such as memory consolidation and recall has been addressed in other computational models (Tetzlaff et al., 2013, 2011; Zenke et al., 2015). Tetzlaff et al. (2013) proposed a dynamic neural circuit where a combination of Hebbian plasticity and synaptic scaling determines memory maintenance during synaptic reactivation. In this setup, memories with weights that surpass a certain threshold are consolidated by spontaneous reactivation, while others decay over time. Scaling in this model also accounted for the destabilization effects of new learning after recall of a motor memory (Walker et al., 2003): if retrieval was followed by learning of a partially overlapping pattern, the weights underlying the first memory were disrupted. This effect is analogous to what was found in our second model, but was observed in the absence of Hebbian plasticity blockade.

### Molecular mechanisms involved in synaptic downscaling and labilization

Although homeostatic plasticity provides an elegant explanation for the dependence of destabilization on memory recall, this is still a speculative hypothesis. We thus attempted to investigate whether there is experimental evidence to support this connection by systematically reviewing the literature. We found that several molecular mechanisms are involved in both memory destabilization and synaptic scaling, suggesting shared requirements for the induction of both phenomena. Studies of destabilization generally test whether pharmacological agents can prevent the reconsolidation-blocking effects of drugs such as protein synthesis inhibitors in behavioral tasks; thus, detecting relevant studies is relatively straightforward. The literature on homeostatic plasticity, on the other hand, is more varied, using different models to study synaptic responses to changes in firing rate and/or network activity. Thus, it is not always clear whether phenomena are comparable across diverse models and preparations, even within the specific field of synaptic downscaling.

Investigations of labilization mechanisms typically use PSIs such as anisomycin to block reconsolidation, indicating that they do not prevent memory destabilization Thus, a requirement for a putative synaptic mechanism of destabilization is that it should not critically depend on protein synthesis – or at least should be less reliant on it than Hebbian plasticity. For synaptic downscaling, different effects of PSIs have been observed. Goold and Nicoll (2010) demonstrated that cycloheximide and anisomycin blocked the AMPAR, but not the NMDAR component of synaptic downscaling after chronic optogenetic stimulation. On the other hand, Hou et al. (2011) showed that anisomycin did not interfere with GluA1 reduction induced by light-gated GluR6 stimulation in hippocampal cultures. Additionally, a recent study performed a comprehensive mapping of synaptic proteins regulated by synaptic scaling, showing that most synaptic proteins exhibited a decrease in synthesis during bicuculline-induced downscaling (Dörrbaum et al., 2020). Thus, the requirement of protein synthesis for downscaling seems to differ across models; nevertheless, at least in some instances, this type of plasticity seems to occur in spite of protein synthesis inhibition.

The majority of mechanisms implicated in destabilization have been identified in studies of fear conditioning, and concern signaling via cell surface receptors. These include GluN2B-containing NMDARs (Ben Mamou et al., 2006; Merlo et al., 2015; Nakayama et al., 2016; Yu et al., 2016) and LVGCCs (De Oliveira Alvares et al., 2013; Haubrich et al., 2015; Kim et al., 2011; Suzuki et al., 2008), as well as, the internalization of GluA2-containing AMPARs (Hong et al., 2013; Shehata et al., 2018). There is evidence that each of these cell surface mechanisms is important for synaptic downscaling. NMDARs are necessary for synaptic downscaling induced by bicuculline in hippocampal slice cultures (Qiu et al., 2012) and dissociated hippocampal neurons (Lee and Chung, 2014), although Goold and Nicoll (2010) showed that APV treatment did not inhibit optogenetically-induced downscaling in hippocampal cultures, suggesting that the role of NMDARs is associated with glutamatergic-driven network activation. AMPARs are a primary mechanism for expression of homeostatic plasticity through alterations in their number, composition, and biophysical properties (Diering and Huganir, 2018). Supporting evidence includes the observations that synaptic scaling relies on a switch between GluA2-containing and GluA2-lacking AMPARs (Chowdhury and Hell, 2018), and that optogenetically-induced AMPAR downscaling does not occur in GluA2-deficient mice (Goold and Nicoll, 2010). Finally, LVGCCs have been shown to be necessary for AMPAR and NMDAR downscaling, both after optogenetic stimulation (Goold and Nicoll, 2010) and after prolonged stimulation with AMPA (Lin et al., 2000) or bicuculline (Lee and Chung, 2014).

Intracellularly, the primary mechanism of destabilization is protein degradation by the UPS, as evidenced by the effect of the proteasome inhibitor lactacystin on destabilization when injected into the hippocampus (Choi et al., 2010; Da Silva et al., 2013; Lee, 2008; Lee and Chung, 2014; Lee et al., 2008), amygdala (Fukushima et al., 2014; Jarome et al., 2011), perirhinal cortex (Stiver et al., 2017) or nucleus accumbens (Ren et al., 2013). Lactacystin also blocks slow homeostatic changes caused by chronic treatment with bicuculline in cultured hippocampal neurons (Jakawich et al., 2010). However, most of the evidence on UPS involvement in synaptic downscaling comes from work with Nedd4 (E3 ubiquitin ligase enzyme), which targets proteins for ubiquitination. Hou et al (2011) demonstrated that Nedd4 expression increases after chronic neuronal stimulation to mediate AMPAR ubiquitination. Other studies showed the involvement of Nedd4 in synaptic downscaling by knocking down its endogenous protein levels (Scudder et al., 2014) and by expressing Nedd4-2, an ubiquitin ligase from the same family, to restore synaptic downscaling (Lee et al., 2018).

A synaptic model for memory destabilization proposed by Finnie and Nader (2012) proposes that reactivation allows calcium influx at the synapse through LVGCCs, with second messengers such as protein phosphatases altering neuronal excitability by modulating NMDARs and AMPARs at the synapse through processes such as GluA2 endocytosis. One would expect that, if this hypothesis holds true, the calcium influx promoted by LVGCCs during memory reactivation might trigger synaptic downscaling pathways that use molecules such as Arc/Arg3 (Chowdhury et al., 2006) and MeCP2 (Qiu et al., 2012), known to regulate AMPA receptor trafficking in models of homeostatic plasticity.

### Theoretical and experimental evidence for a role of homeostatic plasticity in destabilization

Theories postulating a role for homeostatic plasticity in memory need to take into account that homeostatic processes such as synaptic scaling operate in a timescale of hours or days (Zenke and Gerstner, 2017). Thus, the latency for a memory to decay after reconsolidation blockade – around 4 hours after protein synthesis blockade (Nader et al., 2000) – could be related to the time required by homeostatic plasticity to occur (Ibata et al., 2008). One criticism of this assumption is that synaptic scaling is usually induced by hours of continued overstimulation, something that is unlikely to occur during or after memory retrieval *in vivo*. However, Mendez et al. (2018) showed changes reminiscent of homeostatic plasticity after 10 minutes of *in vivo* stimulation, while Moulin et al. (2019) observed similar effects with low-frequency stimulation over a 24h period. Thus, it is possible that *in vivo* homeostatic plasticity could result from periodic reactivations of memory engrams occurring during the post-reexposure period (Wittenberg et al., 2002).

The idea that homeostatic plasticity can play a role in memory processes is not new; however, it has not been explored in detail, as shown by a systematic review of the literature linking both processes (see Methods and **Table S2** for full results). While there are computational models suggesting that homeostatic plasticity is related to memory phenomena (de Camargo et al., 2018; Susman et al., 2019; Tetzlaff et al., 2013), experimental evidence for this relationship is still scarce. Perhaps the most direct attempt at connecting both processes comes from Mendez et al (2018), who used optogenetically-induced spike trains in hippocampal granule cells of mice to trigger *in vivo* homeostatic changes. This protocol decreased excitatory synaptic density while increasing inhibitory synapses through an LVGCC-dependent mechanism. When these spike trains were induced during an extinction session, lower freezing activity was observed in a subsequent test. These results are in line with the view that, at least in some cases, extinction might have a destabilization component occurring along with the learning of a new association, as suggested by previous work (Almeida-Corrêa and Amaral, 2014; Barad, 2006; Popik et al., 2020) and by our own model results (see **Figure 2d**).

Another set of views postulating a role for basic plasticity mechanisms in memory destabilization comes from the observation that reconsolidation-like effects are not restricted to learning paradigms in vertebrates, but are also observed in more reductionist preparations. Studies of the gill-and-siphon withdrawal reflex in Aplysia showed that non-associative long-term facilitation was reversed when heterosynaptic reactivation occurred in the presence of rapamycin, an inhibitor of ribosomal protein synthesis (Hu and Schacher, 2014). For associative long-term facilitation, on the other hand, destabilization required homosynaptic reactivation (Hu and Schacher, 2015). Reconsolidation-like phenomena were also observed in spinal cord pain processing circuits of mice injected with capsaicin to induce mechanical hyperalgesia, with anisomycin reducing hyperalgesia only when paired with another capsaicin injection. (Bonin and De Koninck, 2014) This prompted the authors to postulate that such effects could be due to homeostatic plasticity mechanisms, and that reconsolidation might not be best conceptualized as a behavioral phenomenon (Bonin and De Koninck, 2015).

### Conceptual gaps and future directions

Although the destabilization-reconsolidation process has typically been hypothesized to serve a high-level cognitive function in memory updating (Exton-McGuinness et al., 2015; Fernández et al., 2016; Lee, 2009), it is important to recognize that its existence does not imply a functional role (Dudai, 2004). Nevertheless, the accumulation of evidence that the destabilization of memories occurs preferentially under conditions of memory updating (Lee, 2009; Rodriguez-Ortiz and Bermúdez-Rattoni, 2017) strongly supports a cognitively functional process. This contrasts with the more basic compensatory nature of homeostatic plasticity (Siddoway et al., 2014), which is usually seen as a low-level property of neuronal physiology. That said, categorization of mechanisms as high-level/cognitive vs low-level/physiological may represent an artificial dichotomy, as evolution can lead to repurposing of traits or mechanisms for different purposes than those for which they have evolved (Lloyd and Gould, 2017).

The fact that many cellular mechanisms that are functionally implicated in destabilization have also been shown to be important for synaptic downscaling (at least under certain circumstances) opens up the possibility that destabilization might be an emergent property of homeostatic plasticity that arises from patterns of neuronal activity induced by memory reactivation. Moreover, if homeostasis is detrimental to the previously adjusted synaptic weights that encode a memory in a neuronal network, this would justify the requirement of a reconsolidation process to ensure preservation of the memory trace. This adds to the argument that destabilization-reconsolidation is a universal property of memories (Lee, 2009) as in this case it would emerge from a fundamental property of neuronal function.

The results of our second model are consistent with downscaling being preferentially engaged under conditions of conflicting information or mismatch during memory retrieval that would be expected under memory-updating conditions. In this simple attractor network, this happens as a natural consequence of noisier retrieval when patterns diverge between training and reexposure. Such a view must be balanced, however, against the evidence for the involvement of prediction error signals in destabilization (Das et al., 2015; Exton-McGuinness et al., 2015; Reichelt et al., 2013; Sinclair and Barense, 2019). Detection of mismatch/prediction error within memory-encoding areas could thus be implemented both by novelty signals sent by other brain structures (such as the ventral tegmental area) and by internal network dynamics (as occurs in our model). Support for this view comes from the asymmetry in the necessity and sufficiency of destabilization mechanisms: while dopamine D1 receptors are necessary for fear memory destabilization, for instance (Flavell and Lee, 2019; Merlo et al., 2015), their activation is not sufficient to induce it (Flavell and Lee, 2019). Thus, it is possible that multiple processes are necessary for successful memory destabilization, including an interaction between dopaminergic signaling and local network plasticity.

Many other questions still need to be addressed before a role for homeostatic plasticity in memory destabilization can be asserted. Classic destabilization studies with reversal of reconsolidation blockade could be used to study whether canonical molecular components of homeostatic plasticity, such as the immediate-early genes Homer1a (Hu et al., 2010) and Arc/Arg3.1 (Gao et al., 2010) are involved in destabilization. Another molecule involved in synaptic scaling that has not been implicated in destabilization is brain-derived neurotrophic factor (BDNF) (Reimers et al., 2014). Interestingly, even though BDNF has been extensively implicated in memory consolidation (Bekinschtein et al., 2014), it has been suggested not to be as important in reconsolidation (Lee, 2008; Lee et al., 2004). This could be due to the fact that, if inhibiting BNDF function affects not only Hebbian plasticity but destabilization mechanisms as well, it could lead the net effect of this intervention on a reactivated memory to be neutral. Nevertheless, evidence obtained by investigating the role of synaptic scaling mechanisms in memory destabilization paradigms will still be correlative. For a causal link, experiments that can induce or inhibit homeostatic plasticity in neurons related to a fear memory engram through artificial stimulation can provide more direct evidence. Particularly, if methods for optogenetically-induced homeostatic plasticity *in vivo* such as that used by Mendez et al. (2018) indeed trigger destabilization, pairing this stimulus with reconsolidation blockers such as anisomycin should lead to a reduced fear response in animals.

Even with these more sophisticated studies, however, one still runs into the question of specificity – after all, it is unlikely that the effects of neuronal overstimulation are limited to a particular plasticity process (Keck et al., 2017). Nevertheless, the notion of a ‘particular plasticity process’ might be in itself a paradigm in need of revision, as the division between well-delineated, individual classes of plasticity is more of an epistemic convention than a fact of nature. With a plethora of processes in distinct models and preparations falling within the umbrella of homeostatic plasticity, the most useful way to classify them in order to advance research is not obvious, as both overgeneralization and overspecificity can hamper progress (Fox and Stryker, 2017). Combining advances in this field with those in memory research and its own distinct paradigms presents a further challenge that can only be overcome by better communication between experimentalists and theorists on both sides.

## MATERIALS AND METHODS

### Computational models

#### Model 1 – Adaptation of Osan et al. (2011)

The network consists of a circuit with 100 units connected in an all-to-all manner. Each neuron *i* in the attractor network has a neuronal activity *u_i_* which varies continuously from 0 to 1 and changes according to

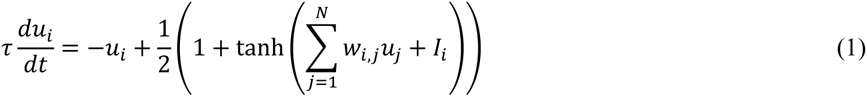

where τ is the neural time constant, which is a combination of properties that denote decay or persistent activity outside the task, and *w_i,j_* represents the synaptic weight associated with a particular connection. *I_i_* represents the input provided to the memory network by sensory stimulus from the environment and internal information. The feedback between the cue network and memory area is not explicitly modeled; nevertheless, it is assumed to be necessary to account for changes in the animal’s internal representation according to learned experience.

Weight changes (ΔW) in the model can be described as

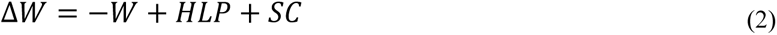

where HLP is the Hebbian learning plasticity term and SC is the synaptic scaling term. Both terms are matrices that are dependent on neuronal activation reached after cue presentation.

The Hebbian learning plasticity term can be described as

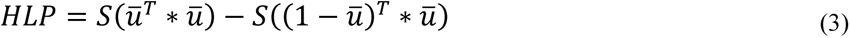

where *S* is a factor that represents the biochemical requirements of Hebbian plasticity and 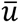 is a vector (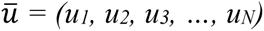) that represents the stable state of network activity. Thus, when two neurons fire together, their connections are reinforced. If a presynaptic neuron fires while the postsynaptic one is inactive, an inhibitory connection is created. Although a realistic implementation of the rule would require modeling the participation of inhibitory interneurons in the process, this simplification allows attractor functioning to occur while eliminating the artificial negative activations and ‘mirror attractors’ found in the original Hopfield graded activity formulation (Hopfield, 1984).

The synaptic scaling term can be described as

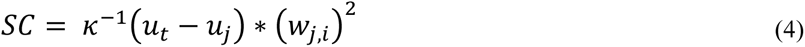

where *κ* is the ratio between Hebbian plasticity and scaling and *u_t_* is the desired homeostatic activity for the network. Therefore, synaptic scaling adjusts connection weights on the basis of a comparison between a neuron’s output activity *u_i_* against a desired target activity *u_t_*. The term is only active if the connection between the pre- and post-synaptic neuron is excitatory.

Learning is induced by providing a noisy input equals to 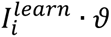 for neuronal activation and –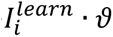 for neuronal inhibition. The input is modulated by a noise term *ϑ* from a uniform distribution [0.9, 1.1]. In the training session, the neurons representing context and aversive neurons are activated while other neurons are inhibited. In the nonreinforced reexposure session, the cue input is given by

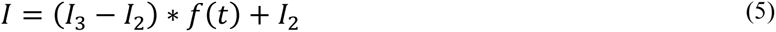

where vectors *I_2_* and *I_3_* represent the cue inputs for the training and extinction pattern, respectively, and *t* represents reexposure duration, varying from *t_min_* = 0 to *t_max_* = 10. For the encoding of reexposure time in the input cues, we use a sigmoid function defined as:

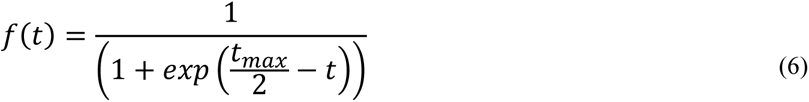

Retrieval tests consist of activating the neurons that represent the context while maintaining the input to other units at 0. After the input is given, the pattern to which the network evolves determines the degree of freezing in the test session, with retrieval of the training pattern resulting in 90% freezing and other patterns resulting in 10% freezing. For pattern determination, we compare the current activity of each neuron with that of the same neuron in the training pattern. If a neuron’s output activity *u_i_* is greater than 0, the neuron is considered to be active; otherwise, is considered to be silent. The fear memory is considered to be successfully retrieved when it has more than 95% similarity with the training pattern in this binary classification. We perform 100 simulations with different initial conditions in the retrieval session and obtain the mean ± S.E.M. freezing behavior for each reexposure time. All model simulations are developed using Matlab R2018a.

To check the robustness of our results to different parameters, we vary the value of S during training or reexposure and the value of κ during reexposure. S during reexposure is varied from 0 to 1 with steps of 0.025 in Fig. S1. κ during reexposure is varied from 0 to 4 with steps of 0.2 in Fig. S2. S during training is varied from 0 to 2 with steps of 0.1 in Fig S3.

If not stated otherwise, we used the parameters described in **Table 4**. Code for the simulations is available at https://github.com/Felippe-espinelli/scaling_destabilization_models.

**Table 4.** Model parameters used by the attractor network adapted from Osan et al (2011).

| Variable | Description | Value |
| --- | --- | --- |
| $S$ | Biochemical requirements for Hebbian synaptic plasticity | 0.8 |
| $u_t$ | Desired target activity of synaptic scaling | 0 |
| $\kappa$ | Ratio between Hebbian plasticity and scaling | 1.2 |
| $N$ | Number of neurons | 100 |
| $\tau$ | Neural time constant | 1 |
| $I_i^{learn}$ | Learning input strength | 8 |
| $I_i^{ret}$ | Retrieval input strength | 8 |

#### Model 2 – Adaptation of Auth et al. (2020)

The model is composed of an input network and a memory network with 36 and 900 neurons, respectively. The memory network is a grid of 30 x 30 neurons organized in a toroidal topology; each of them connects with 4 random neurons in the input area and with their nearest-neighbors in the memory area within a radius of 4 neurons. An inhibitory unit represents a population of inhibitory neurons connected bidirectionally in an all-to-all manner with neurons in the memory area.

For each excitatory neuron *j* in the memory area (*j ∊* {1, …, *N^M^*}), the membrane potential *u_j_* is determined by the following differential equation:

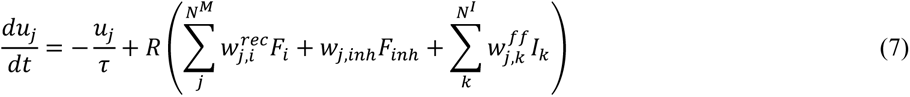

Afterwards, the firing rate *F_j_* of each neuron *j* is dependent of the membrane potential *u_j_* as follows:

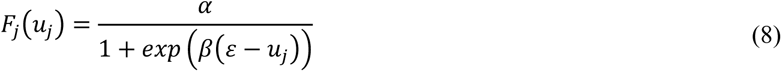

Initial membrane potential and firing rate are drawn from a normal random distribution with a mean of 0 and a variance of 1.

The inhibitory unit updates the membrane potential *u_inh_* and converts it into a firing rate *F_inh_* as described below:

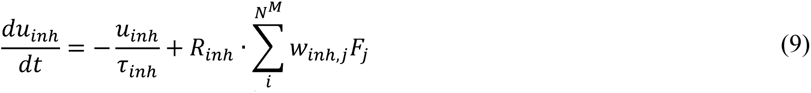

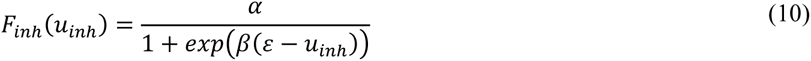

Excitatory feed-forward and recurrent connections are plastic, while others remain constant. These weight changes are dependent on a combination of Hebbian synaptic plasticity and synaptic scaling. The feed-forward synaptic weight 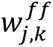, formed by connecting an excitatory input neuron *k* (*k ∊* {1, …, *N^I^*}) to a memory neuron *j* (*j ∊* {1, …, *N^M^*}), is initially drawn from a uniform distribution 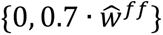 with weight changes described as:

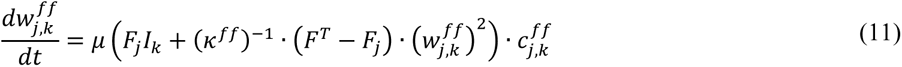

where 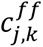 is a feed-forward connectivity matrix, with each unit equal to 1 if the connection exists or 0 if it does not.

Recurrent synaptic weights 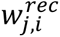, formed by connecting an excitatory memory neuron *i* (*i ∊* {1, …, *N^M^*}) to a memory neuron *j* (*j ∊* {1, …, *N^M^*}) are initially set as 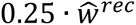 with weight changes described as

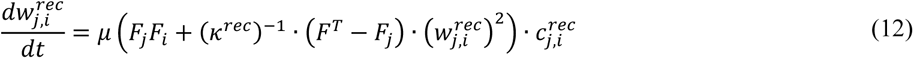

where 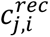 is a feed-forward connectivity matrix, with each unit equal to 1 if the connection exists or 0 if it does not.

A training session is simulated by activating 18 input area neurons with an input rate of 130 Hz, while the remaining ones receive 0 Hz. This activation is performed in 10 periods of 5 s each, with a 1 s rest period between them. For non-reinforced reexposure, we also use 10 periods with 5 s of activation and 1 s of rest. For the shortest possible reactivation (t=0), the input pattern in the reexposure session is equal to the training cue. Progressively longer reexposure durations are simulated by substituting input neurons from the training cue by those representing the extinction pattern, two at a time. Thus, the longest possible reexposure (t=18) corresponds to the full extinction pattern, while for intermediate values t corresponds to the number of divergent neurons between the training and reexposure patterns. As in the training session, activated neurons are stimulated at 130 Hz while the others are set at 0 Hz.

All model simulations were developed using Python 3.7, with the code available at https://github.com/Felippe-espinelli/scaling_destabilization_models. Equations were solved using the Euler method with a time step of 0.005 s. For each time step, the network is updated in the following order, (1) membrane potential of each neuron in memory area; (2) membrane potential of inhibitory unit; (3) firing rate of memory area; (4) firing rate of inhibitory unit; (5) recurrent weight; (6) feed-forward weight. All model parameters are described in **Table 5**.

**Table 5.** Model parameters used in the adaptation of Auth et al. (2020).

| Variable | Description | Value |
| --- | --- | --- |
| $\tau$ | Membrane time constant (memory area) | 0.01 |
| $R$ | Membrane resistance (memory area) | 1/11 |
| $N^M$ | Number of neurons in memory area | 900 |
| $N^I$ | Number of neurons in input area | 36 |
| $I_k$ | Input rate | {0,130} |
| $\alpha$ | Maximum firing rate | 100 |
| $\beta$ | Sigmoid steepness | 0.05 |
| $\varepsilon$ | Sigmoid inflexion point | 130 |
| $\mu$ | Plasticity time constant | 1/15 |
| $F^T$ | Target firing rate | 0.1 |
| $\kappa^{rec}$ | Scaling time constant (recurrent) | 60 |
| $\kappa^{ff}$ | Scaling time constant (feed-forward) | 720 |
| $\tau_{inh}$ | Membrane time constant (inhibitory unit) | 0.02 |
| $R_{inh}$ | Membrane resistance (inhibitory unit) | 1 |
| $w_{inh,i}$ | Synaptic weight: memory area to inhibitory unit | 0.6 |
| $w_{j,inh}$ | Synaptic weight: inhibitory unit to memory area | 6000 |
| $\hat{w}^{rec}$ | Initial recurrent synaptic weights | $\sqrt{\frac{\kappa^{rec} \alpha^2}{\alpha - F^T}}$ |
| $\hat{w}^{ff}$ | Initial feed-forward synaptic weights | $\sqrt{\frac{\kappa^{ff} \alpha \cdot 130}{\alpha - F^T}}$ |

#### Cluster analysis

Each consolidated memory is represented by the network as a cluster of neurons recurrently connected by strengthened memory weights. Memory clusters after the training session are defined as neurons with a mean postsynaptic weight for all their recurrent connections above a threshold *ϕ*, set to 40 for the main simulations. As random connections can occasionally rise above this threshold, we only define a cluster if a minimum amount ρ of 30 neurons is above the threshold. When a cluster is identified, the mean connection weight between neurons belonging to the cluster is used as a measure of memory strength. If no clusters are identified, memory strength is defined as the global mean weight of the network.

To identify modifications in consolidated clusters and formation of new ones after reexposure, we initially identify the training session cluster. Neurons that have a recurrent connection with any neuron from the original training cluster are considered to be part of the training cluster if the mean weight of all their postsynaptic connections is above the threshold *ϕ*. This allows slight modifications in cluster position to be viewed as an update of the original memory rather than a new cluster. Neurons with recurrent weights above the threshold that do not connect to the training cluster will be considered as being part of a new (extinction) cluster. Note that all active neurons that do not belong to the training cluster will be placed in the extinction cluster, even if they have no mutual connections between them.

### Systematic review of labilization mechanisms in different structures

#### Search strategy

The protocol for this systematic review was preregistered in the Open Science Framework at 10.17605/OSF.IO/ZHPR4. Briefly, we performed a search in PubMed and Web of Science using the search terms *(destabili* OR labil*) AND (reconsolidat* OR reactivat*)*, including studies published until November 4^th^, 2019. These terms were developed and refined based on a systematic review performed by our group as part of a previous study (Lee et al., 2019)

#### Study selection

Two independent investigators (F.E.A. and R.L.C.) screened titles and abstracts for (i) original studies; (ii) written in English, (iii) that included experiments evaluating the modulation of reconsolidation blockade by an intervention targeting a specific molecular mechanism. We used the Rayyan platform (Ouzzani et al., 2016) to select studies and exclude duplicates. An article proceeded to the full-text screening stage if it was included by at least one reviewer. Agreement between investigators was 98.3%.

Inclusion criteria in full-text screening (which included supplemental material when available) were the following: (i) studies describing the behavioral effects of an intervention directed at a specific molecular mechanism, (ii) performed up to 6h before or after a reactivation session (iii) that modulated the effect of reconsolidation blockade by another drug (e.g., anisomycin and MK-801). As in the previous step, two reviewers (F.E.A. and R.L.C.) evaluated all studies. Disagreements were discussed and solved with the help of a third investigator (O.B.A.).

Information from each article was extracted by a single reviewer and reviewed by the other. Extracted variables included the reconsolidation inhibitor used, its injection time and site, the destabilization treatment with its own infusion time and site, the molecular target of the treatment and the behavioral outcome as described by the authors, All extracted information was inserted in a .xls spreadsheet (**Supplementary Raw Data 1**).

### Review of the role of memory destabilization mechanisms in synaptic downscaling

To search for articles relating mechanisms of memory destabilization to synaptic downscaling, we initially analyzed the dataset from a recent scoping review of the field that included 168 studies (Moulin et al., 2020). One of the authors (T.C.M) screened the full text of 51 articles in which chronic excitation experiments were performed to induce downscaling, along with pharmacological or genetic interventions to investigate the underlying molecular machinery. All articles reporting results relating synaptic downscaling to one of the memory destabilization components described in Table 2 were selected for further analysis, irrespective of their results.

As the analyzed systematic review dataset comprised studies published until 2017, we manually performed individual searches combining the term “synaptic scaling” with keywords associated with the molecular mechanisms shown in table 2 to obtain updated results. The searches were performed on June 29^th^, 2020 using the following search terms: “synaptic scaling AND NMDA” for NMDA receptors; “synaptic scaling AND (UPS OR ubiquitin*)” for the ubiquitin-proteasome system; “synaptic scaling AND AMPA” for AMPA receptors; “synaptic scaling AND calcineurin” for calcineurin; “synaptic scaling AND L-type calcium channel” for LVGCCs; “synaptic scaling AND (PP1 OR protein phosphatase-1)” for PP1; “synaptic scaling AND CaMKII” for CaMKII; “synaptic scaling AND (sodium channels OR Na channels)” for Na^+^ channels; “synaptic scaling AND (“protein synthesis” OR cycloheximide OR anisomycin)” for protein synthesis; “synaptic scaling AND (cholinergic receptor OR acetylcholine receptor)” for cholinergic receptors; “synaptic scaling AND CB1” for CB1 receptors; “synaptic scaling AND nitric oxide” for nitric oxide; “synaptic scaling AND dopamine receptor” for dopaminergic receptors; “synaptic scaling AND autophagy” for autophagy; “synaptic scaling AND histamine receptor” for H3 receptors; “synaptic scaling AND IP3” for IP3Rs; and “synaptic scaling AND beta adrenergic receptor” for β-adrenergic receptors. The term “NOT review” was added to all searches in order to narrow the results to original studies.

Articles investigating synaptic downscaling were identified through abstract and full-text screening by a single investigator (T.C.M.). Inclusion criteria were (i) articles written in English, (ii) presenting original results, and (iii) describing animal experiments using chronic stimulation of neurons to study synaptic downscaling. If the title and abstract were not clear about the three criteria described above, articles were still considered for full-text screening. Articles were included after full-text screening if they described experiments manipulating and/or measuring the activity or expression of the molecule of interest as part of a protocol previously or concurrently shown to induce synaptic downscaling. The main findings and brief descriptions of the methods and biological models from the 13 articles describing the involvement of a memory destabilization mechanism in synaptic downscaling were included in Table 3.

### Systematic review of connections between homeostatic plasticity and memory phenomena

#### Search strategy

A systematic review was performed in PubMed and Web of Science using the search terms ((“destabilization” OR “destabilisation” OR “labilization” OR “labilisation” OR “labile” OR “reconsolidation” OR “reactivation” OR “extinction*” OR “recall” OR “retrieval” OR “update” OR “updating” OR forget*) AND (“memory” OR “learning” OR “conditioning” OR “plasticity”) AND (“homeostatic” OR “synaptic scaling” OR “heterosynaptic” OR “metaplasticity”)). We refined search terms in order to include studies that we considered important to the subject prior to the review (Bonin and De Koninck, 2014; Mendez et al., 2018; Tetzlaff et al., 2013, 2011; Zenke et al., 2015). The search included articles published until November 4^th^, 2019, and there were no exclusion criteria for article type.

#### Study selection

Title and abstract screening excluded (i) articles not in English, and (ii) articles without mentions to memory phenomena such as reconsolidation, extinction or labilization or homeostatic plasticity. To be included, at least one reviewer had to include the reference for it to proceed to the next screening stage. Agreement between reviewers in this stage was 84.3%.

The second screening stage considered the full text of the article, including supplemental material if available. Two investigators (F.E.A. and R.L.C.) included studies for later data extraction if they demonstrated or discussed a direct relationship between memory processes (e.g., reconsolidation, extinction, and labilization) and homeostatic plasticity. A reason for the exclusion of an article had to be included in this step. Disagreements between reviewers in this step were solved with the help of a third one (O.B.A.). Included studies were used for the discussion section and are listed in **Table S2**, with the search flowchart presented in **Figure S8**.

## ACKNOWLEDGEMENTS

This work was supported by CNPq scholarships to F.E.A., T.C.M., a PIBIC/UFRJ scholarship to R.L.C. and grant E-26/010.002674/2014 from FAPERJ and the University of Birmingham to O.B.A. and J.L.C.L. The funders had no role in the design, analysis or reporting of the study.

## SUPPLEMENTARY MATERIAL

**Figure S1:**
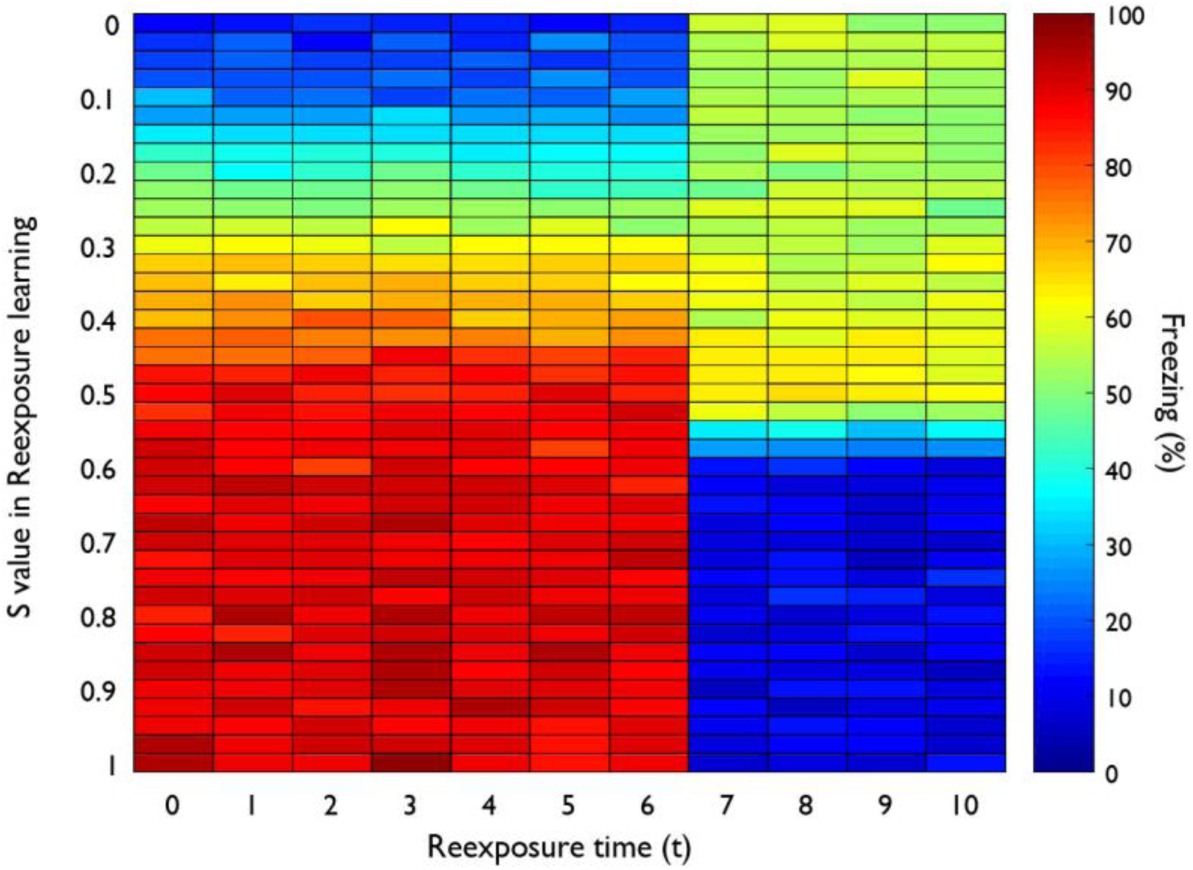
Effects of variable S values during reexposure on test freezing. Heat map shows the effects of variable S values (y axis) during reexposure sessions of various durations (x axis) on mean test freezing (color scale) for 100 simulations. High S values (S > 0.6) lead to a nonlinear transition from reconsolidation to extinction with increasing reexposure time, as observed in the control groups (S=0.8) in Fig. 2. Low S values (S = 0 to 0.15) lead to reconsolidation blockade (blue bins) with lower reexposure times and maintenance of freezing (red bins) with larger reexposure values, as observed in the PSI groups (S=0) in Fig. 2. N = 100 simulations. y-axis: S values vary with steps of 0.025.

**Figure S2:**
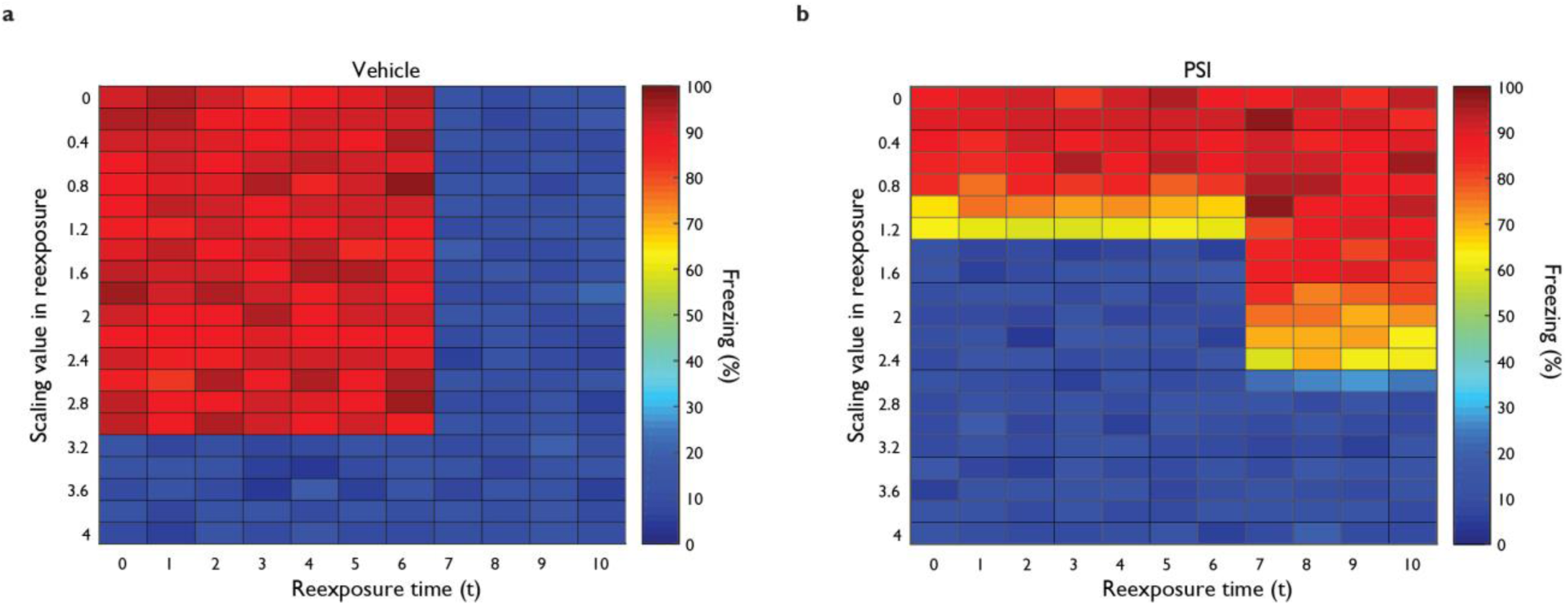
Effects of variable κ values during reexposure on test freezing. **(a)** Heat map shows the effects of varying κ values (y axis) in reexposure sessions of various durations (x axis) on mean test freezing (color scale) in the control group (S = 0.8) for 100 simulations. The nonlinear transition from reconsolidation to extinction still occurs with low values of scaling and only disappears at high values (κ > 3), in which the network is not capable of maintaining previous plasticity in activated neurons. **(b)** Effects of varying κ values during reexposure sessions under Hebbian plasticity blockade (S=0). At lower scaling values, freezing remains high as the training memory is preserved. With higher values, the training memory is degraded with κ values above 1 with t values between 0 and 7. As in the control group, higher values of κ degrade the training memory irrespective of reexposure time. N = 100 simulations. y-axis: Scaling values vary with steps of 0.2.

**Figure S3:**
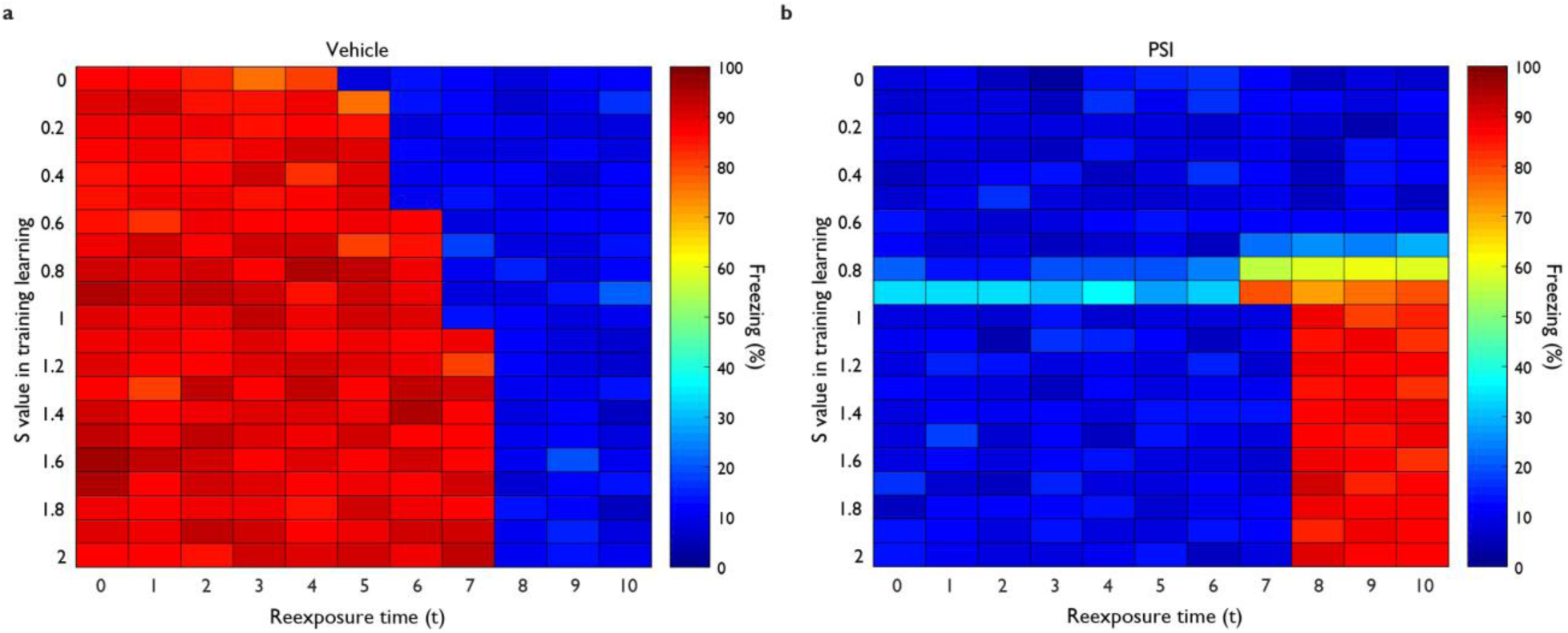
Effects of variable S values during training on test freezing. **(a)** Heat map of the effects of varying S during training (y Axis) for various reexposure times (x axis) on mean test freezing (color scale) in the control group for 100 simulations. Freezing is maintained after shorter reexposures, while extinction occurs at progressively higher reexposure durations as training strength increases. **(b)** Effects of varying S during training for various reexposure times under Hebbian plasticity blockade during reexposure. Freezing is maintained after long reexposures due to extinction blockade for S values o 0.8 or higher, while test freezing is low for all other conditions (blue). N = 100 simulations. S values vary with steps of 0.1.

**Figure S4:**
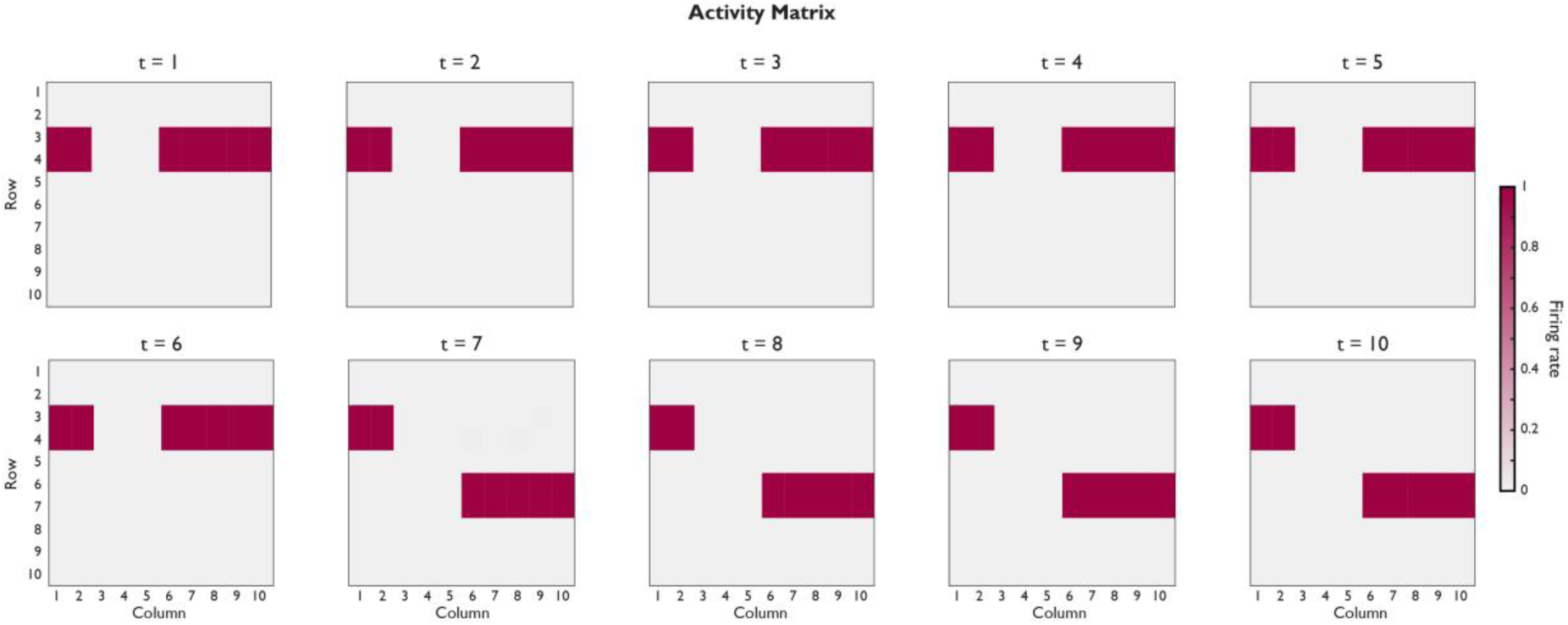
Neuronal activity matrix at the end of reexposure sessions using different reexposure values. Patterns correspond to those shown on Fig. 2, with different neuronal groups representing the context (4 neurons), fear (10 neurons) and safety (10 neurons). With reexposure times between t = 1 and t = 6, the retrieved pattern is identical to that used in the training session, while with longer reexposure durations (t = 7 to t = 10), the network switches to the activity pattern corresponding to the extinction session.

**Figure S5:**
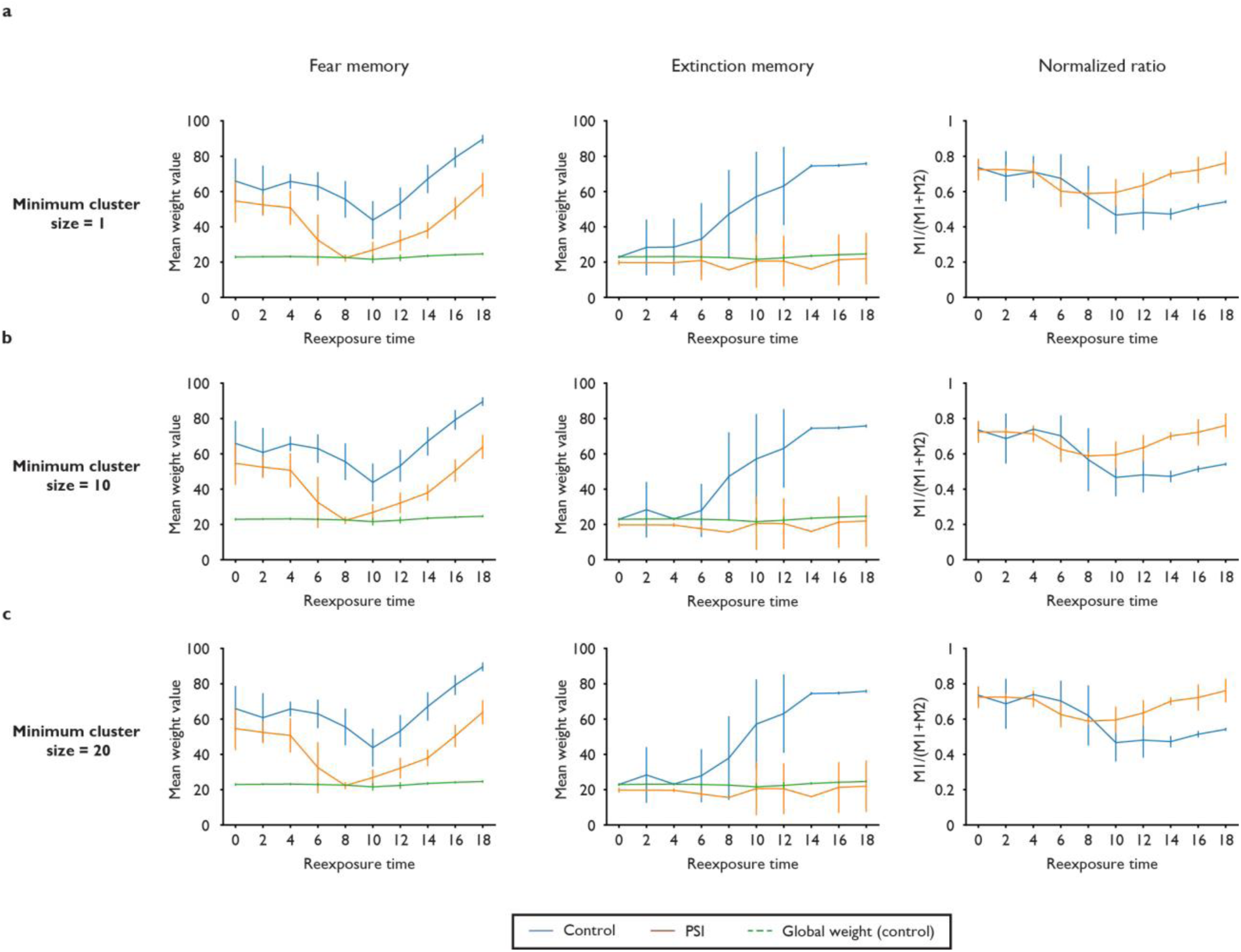
Effects of Hebbian plasticity blockade on synaptic weights using different minimum cluster sizes. Subpanels show the same results as in Fig. 4, but with different minimum numbers of neurons used to define a cluster. **(a)** Recurrent mean weight values for the training memory (left), extinction memory (center) and their normalized ratio (right) under control (blue) and PSI (orange) conditions using a minimum cluster size of 1 neuron. In this case, formation of the extinction cluster occurs at a lower reexposure time compared with the results described in figure 4d. **(b)** Results using a minimum cluster size of 10 neurons. These are similar to the ones using the default values on Fig. 4. **(c)** Results using a minimum cluster size of 20 neurons, also similar to the default values. All results were acquired using the same 20 simulations of Fig. 4, using different criteria for defining clusters.

**Figure S6:**
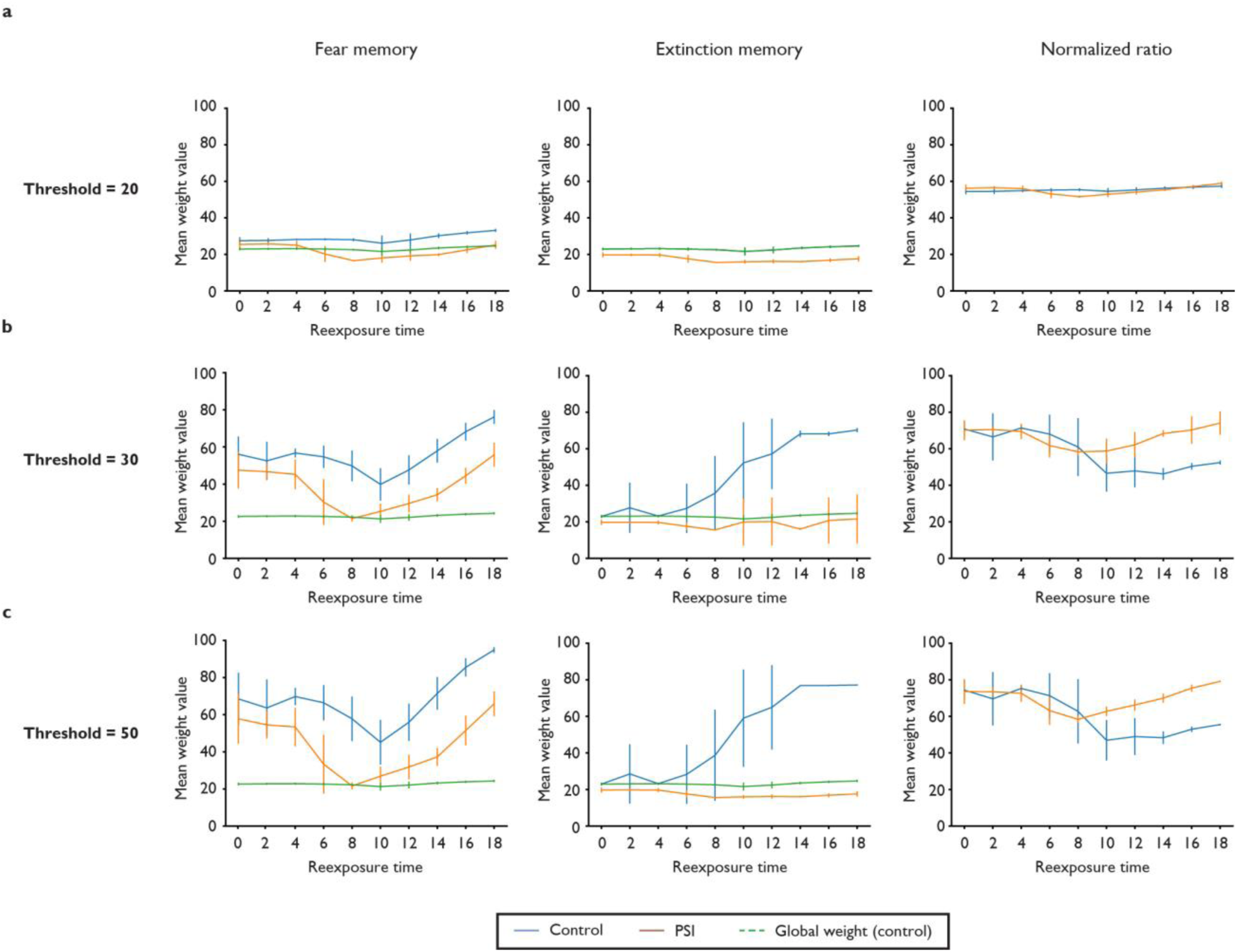
Effects of Hebbian plasticity blockade on synaptic weights using different cluster weight thresholds. Subpanels show the same results as in Fig. 4, but with different mean recurrent weights used to define neurons belonging to a cluster. **(a)** Recurrent mean weight values of the training memory (left), extinction memory (center) and their normalized ratio (right) using clusters defined by neurons with mean weight values above a threshold of 20. With this threshold, clusters have weights that are similar to the global mean weight, indicating that most or all neurons in the memory area are included in the same cluster. **(b)** Results using a weight threshold of 30. These are similar to the ones using the default values on Fig. 4. **(c)** Results using a weight threshold of 50, also similar to the default values. All results were acquired using the same 20 simulations of Fig. 4, using different cluster weight thresholds.

**Figure S7:**
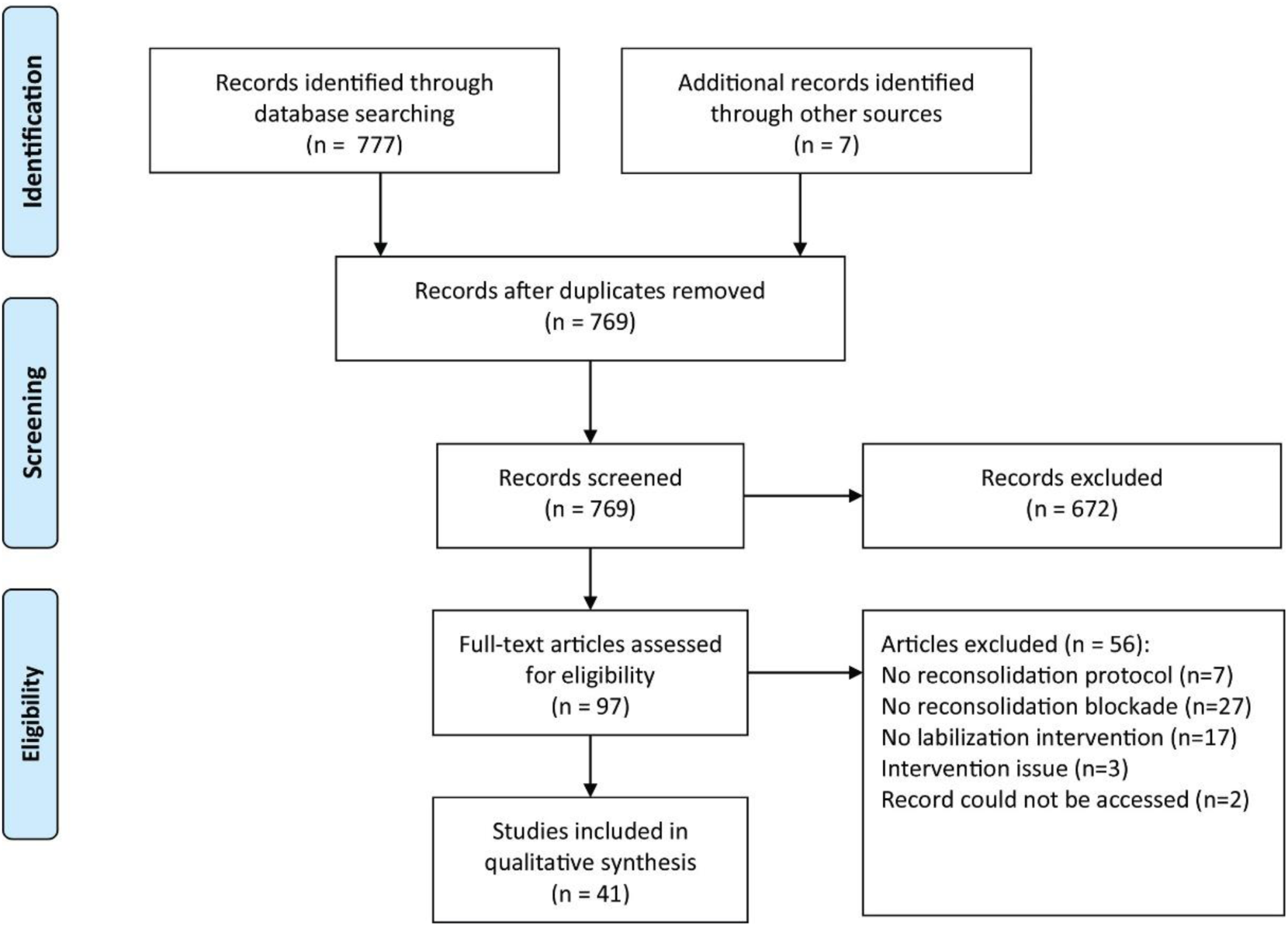
Flowchart of systematic review on labilization mechanisms in different structures. The search process yielded 777 search hits and 7 additional studies were included. With the further exclusion of duplicates, we ended up with 769 articles, of which 672 were excluded during the first screening. For full-text screening, we started with 97 studies, with 41 meeting our criteria for inclusion.

**Figure S8:**
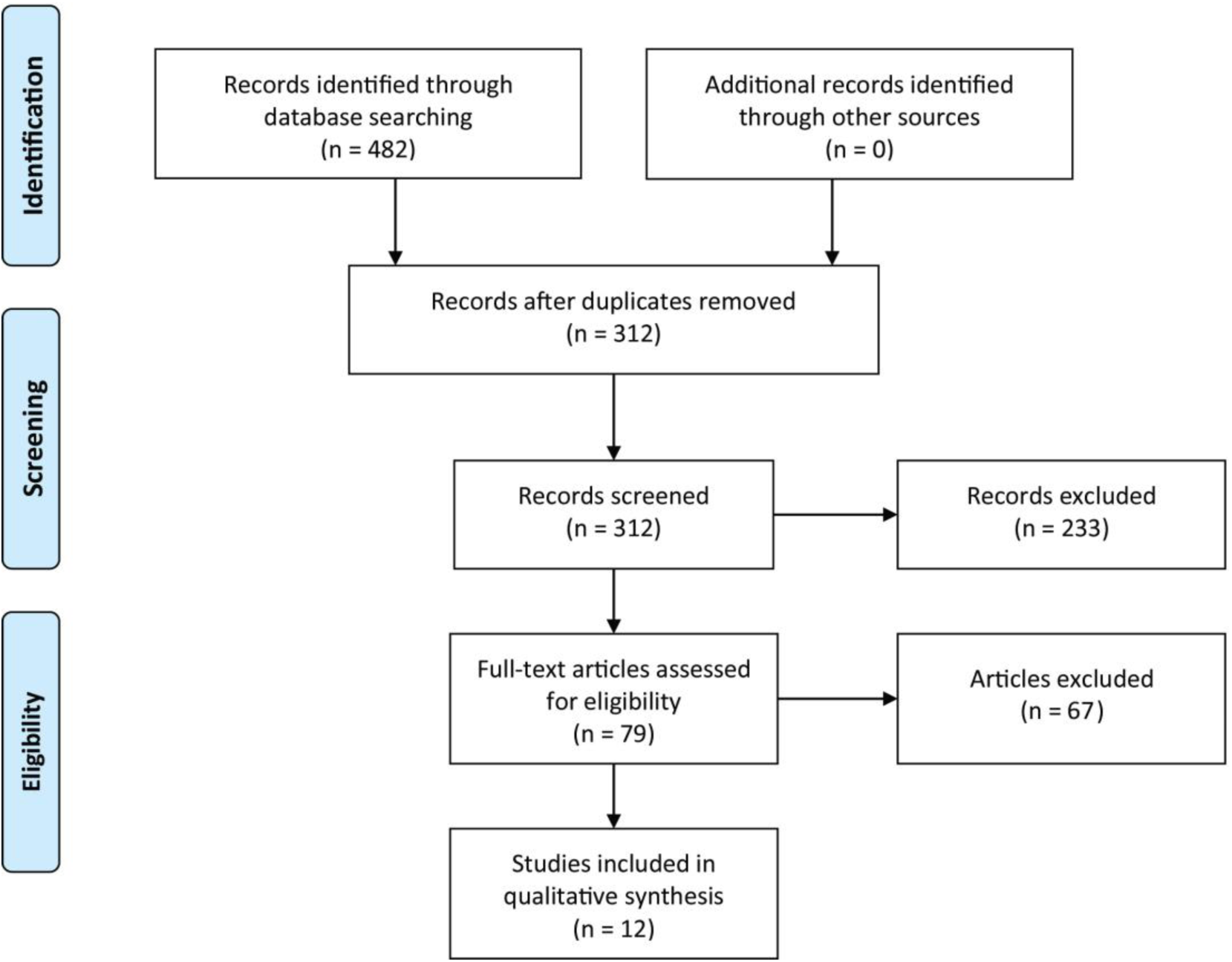
Flowchart of systematic review on relation between homeostatic plasticity in memory phenomena in different structures. The search process yielded 482 search hits. After further exclusion of duplicates, we ended up with 312, of which 233 were excluded during the first screening. In full-text screening, we started with 79 studies, with only 12 meeting our criteria, where 8 were original research articles, 3 were reviews, and 1 was an opinion article.

**Table S1:** Experiments demonstrating boundary conditions of destabilization-blocking interventions. The table shows cases in which negative and positive results were found for the same intervention and brain structure within a single study. For each of these, the boundary condition defining whether results were positive or negative is described. Negative results in which no positive effects of the same intervention were found in a particular brain structure are shown in Table 2. NMDA, N-methyl D-aspartate receptor; CB1, endocannabinoid receptor type 1; I.P., intraperitoneal; H3, histamine H3 receptors.

| Target | # experiments | Boundary condition | Details |
| --- | --- | --- | --- |
| **NMDA** |  |  |  |
| Partial agonism (D-cycloserine) | 1 | Reconsolidation blocker (Ortiz et al., 2015) | An experiment using midazolam as reconsolidation inhibitor showed no effect of D-cycloserine, while another using propranolol showed destabilization enhancement. |
| NR2B antagonism (ifenprodil) | 1 | Timing (Ben Mamou et al., 2006) | Post-reactivation ifenprodil infusion did not prevent reconsolidation blockade, while pre-reactivation ifenprodil infusion blocked destabilization. |
| **CB1 receptors** |  |  |  |
| Antagonism (SR141716) | 2 | Timing (Lee et al., 2019) | In the amygdala, post-reactivation SR141716 infusion did not prevent amnesia induced by MK-801, while pre-reactivation infusion blocked destabilization. In the hippocampus, post-reactivation SR141716 infusion blocked destabilization, while pre-reactivation infusion did not. |
| **β-adrenergic receptors** |  |  |  |
| Antagonism (propranolol) | 1 | Timing (Lim et al., 2018) | I.P. injection of propranolol 30 min before reexposure blocked destabilization. If the injection was performed immediately post-reexposure, there was no effect |
| **H3 receptors** |  |  |  |
| Antagonism (thiopramide) | 1 | Dose (Charlier and Tirelli, 2011) | 5 mg/kg promoted no effect on memory destabilization. Doses of 10 and 20 mg/kg blocked memory destabilization. |

**Table S2:**
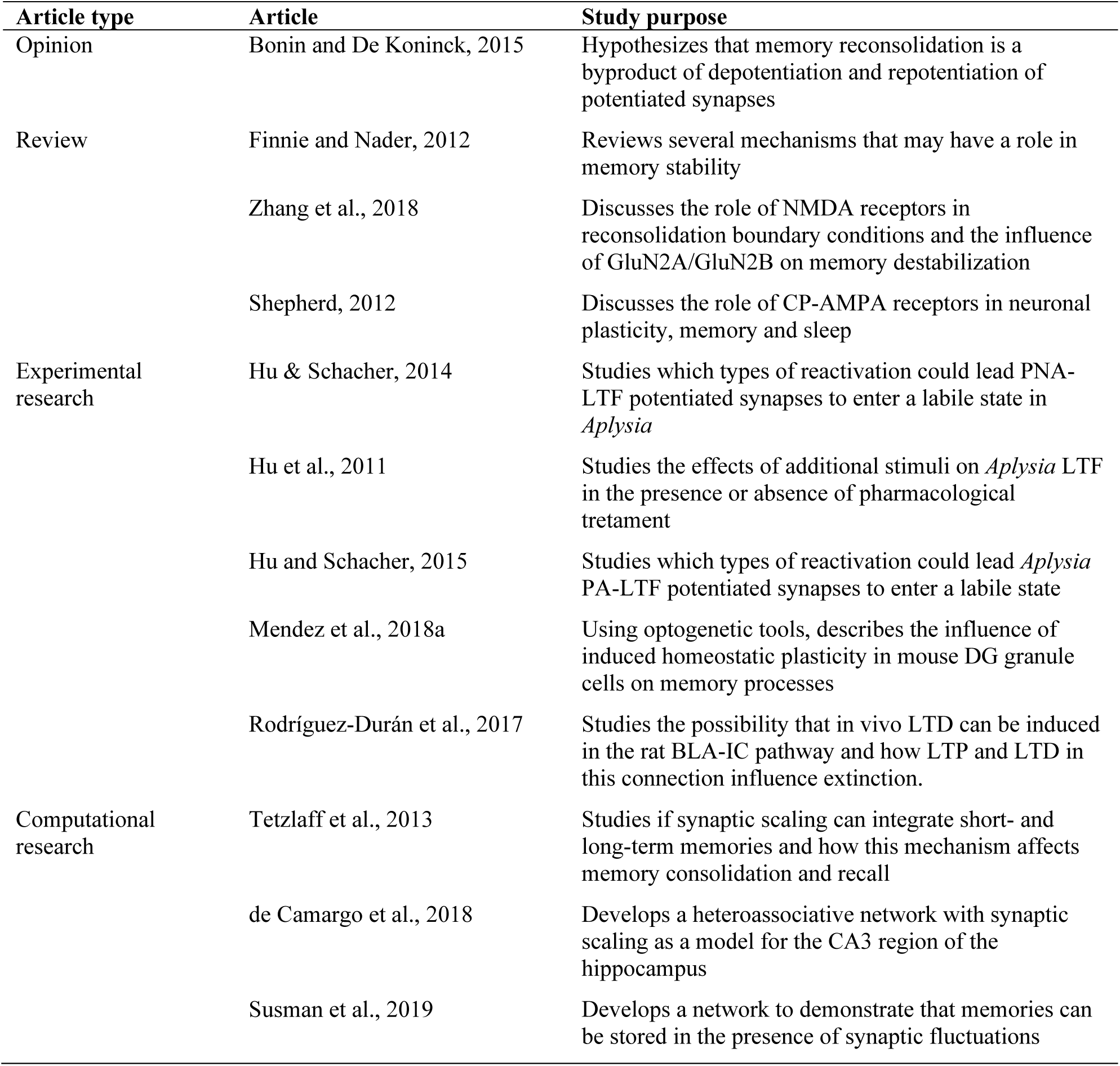
Features of the studies extracted during the systematic review of homeostatic plasticity in memory. Studies are classified as opinion articles, reviews, experimental studies and computational studies according to their methodology. BLA-IC: Basolateral-insular cortex; CP-AMPA: calcium-permeable AMPA; PNA-LTF: persistent non-associative long-term facilitation; DG: Dentate gyrus; LTD: Long-term depression; LTF: Long-term facilitation; LTP: Long-term potentiation; NMDA: N-methyl D-aspartate; PA-LTF: persistent associative long-term facilitation.

## Notes

https://github.com/Felippe-espinelli/scaling_destabilization_models

### Competing Interest Statement

The authors have declared no competing interest.

### Summary of Updates

Acknowledgments updated

